# A Versatile, Simple, and Rapid (VSR) method for generating gene clones, diverse mutants, and short gene fragments for molecular biology applications

**DOI:** 10.1101/2025.10.16.682745

**Authors:** Kavitha Modenkattil Sethumadhavan, Gayatri Kandpal, Jyothilakshmi Sajimon, V. R. Akshay Das, Nirmegh Basu, Athulya Prakash Simla, V. Stalin Raj

## Abstract

Gene cloning, site-directed mutagenesis (SDM), and short gene synthesis are essential tools in molecular biology, yet existing approaches remain constrained by efficiency, flexibility, and cost. Here, we developed a versatile, simple, and rapid (VSR) 3-in-1 method that integrates seamless cloning, mutagenesis, and short gene assembly in a single platform. This approach works on the principles of overlap extension polymerase chain reaction (O_PCR_), uses high-fidelity DNA polymerase and short overlapping primers. Target genes can be directly amplified from genomic DNA or complementary DNA (cDNA) templates and cloned into vectors without intermediate purification, restriction enzyme digestion, or ligation. VSR supports the insertion of DNA fragments ranging from 80 bp to 33 kb at any plasmid locus while enabling the introduction of single or multiple nucleotide substitutions at one or more sites in a single reaction. Using this method, we cloned 35 genes of diverse lengths and two complete viral genomes with efficiencies ranging from 60–100% and introduced up to 17 substitutions in a single mutagenesis reaction. We further demonstrate efficient *de novo* synthesis of short genes from overlapping oligonucleotides. The VSR method is a reliable, rapid, and cost-effective approach with a wide range of applications for both routine and high-throughput workflows.

**GRAPHICAL ABSTRACT:** 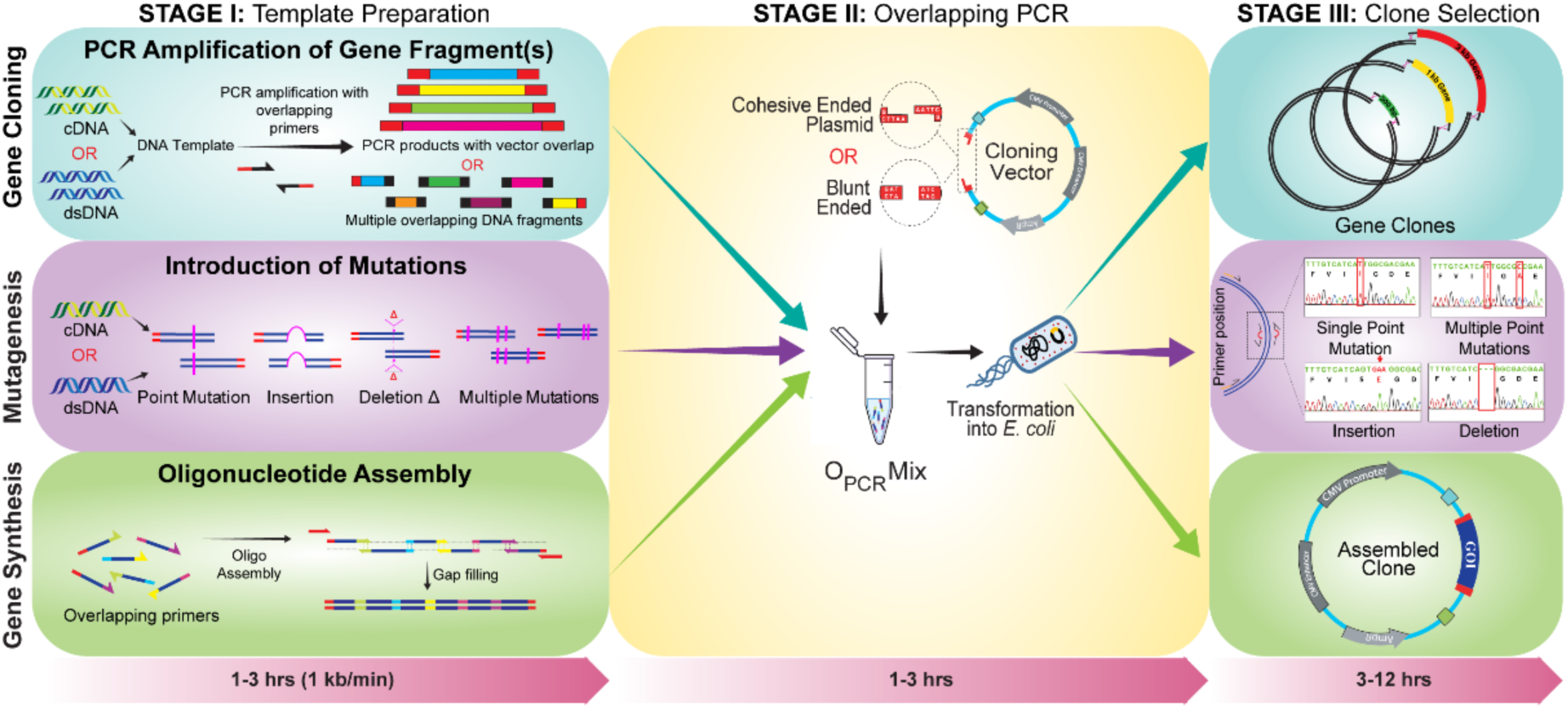

## INTRODUCTION

Gene cloning and site-directed mutagenesis are two fundamental techniques in molecular biology research. These techniques are used to manipulate and identify the role of genes and their encoding proteins. The key steps to evaluate the function of a gene are to clone it into a suitable plasmid vector, mutate through SDM and study the function through *in vitro* studies (1–5). These approaches are fundamental yet critical in a wide range of research fields, including genetics (6), biotechnology (7, 8), microbiology (9), cell biology (10–12), biochemistry (13, 14), genetic engineering (15–17), and ecology (18, 19). Gene cloning and SDM are constantly improved to simplify procedures, reflecting their wide application and the lack of substitutes or alternatives (20, 21).

The first cloning technique developed by Boyer and Cohen is the gold standard in molecular biology (3), but it requires multi-step procedures, which includes the selection of compatible restriction enzymes for the vector and insert, followed by polymerase chain reaction (PCR), restriction enzyme digestion, ligation, bacterial transformation, and screening of colonies, making it a time-consuming approach (22, 23), that often requires several days to weeks to complete the entire protocol. In addition, troubleshooting remains time-consuming and challenging. The recent advancements in recombinant DNA technology has made it possible to develop improved cloning techniques, such as ligation-independent cloning (LIC) (24), Gibson Assembly (25), In-Fusion (26), TOPO (27, 28), sequence and ligation-independent cloning (SLIC) (29, 30), and Gateway cloning (31, 32). But each of these methods has its own pros and cons. For example, Gibson assembly, which is known for its efficiency in assembling and cloning large DNA fragments, but less suitable for cloning short DNA fragments (<250 bp) because of the degradation by the T5 exonuclease present in the reaction mix (33). Similarly, Gateway and TOPO cloning use proprietary commercial vectors (34, 35), which makes it harder to design flexible experiments and raises the overall cost. LIC requires specific 5′ and 3′ overhangs on both the insert and vector, which constrains insert design, whereas improved SLIC is less efficient when the overhangs tend to form stable secondary structures, as it reduces the binding of T4 DNA polymerase (4). Although various cloning methods and kits are available, the ongoing research aims to improve these techniques for efficiency and ease of use. Recently, synthetic plasmid constructs have been used for biotechnological applications to avoid cloning obstacles (36, 37). However, such large-scale gene synthesis or cloning is limited due to higher costs, especially in resource limited laboratory settings (38–40).

Similarly, SDM is a routine and widely used method in molecular biology. Generally, to understand the roles and crucial functions of a gene, amino acids, domains, or motifs in the protein structure, desired mutations are incorporated into the oligonucleotides and introduced in a plasmid through standard PCR amplification (41, 42). As SDM has become an essential tool in the field of molecular biology and genetic engineering, there have been several methods developed in recent years (43–49). However, these approaches have certain limitations, due to the requirement of long complementary primer pairs, which can introduce nonspecific mutations and deletions (50, 51). In addition, the presence of parental plasmid along with the mutants, necessitates sequencing of more clones to obtain desired mutant plasmids (46, 52). Noticeably, while introducing mutations in a large plasmid (>7 kb), the gene of interest is often subcloned into a small vector to improve the mutagenesis efficiency and decrease non-specific mutations. These include multiple steps, including restriction digestion, ligation, transformation, colony selection, DNA isolation, sequence verification, and re-insertion of the mutated sequence into the original plasmid, which is time-consuming and labor-intensive (53). Furthermore, using the conventional methods limits the introduction of multiple or large-scale mutations in a single reaction (1, 54). In contrast to traditional methods such as gene cloning or SDM, gene synthesis involves chemical synthesis of desired genes or plasmids without the need of any template DNA (55, 56). It increases the flexibility in experiments, by allowing customized manipulation of gene sequences, which can result in more advantageous properties than the wild-type sequence (57). Moreover, in situations where clinical materials are difficult to access, for instance, retrieving various sequences of emerging SARS-CoV-2 variants or any genetic disorders, gene synthesis plays a pivotal role in accelerating experimental studies (39, 58, 59). However, synthesizing each variant as an individual construct is not cost-effective, and selectively amplifying minor variants from a mixed population is technically difficult. Therefore, a robust method for cloning and introducing mutations into the backbone of the parental strain would be highly beneficial. Such an approach would allow the systematic generation of all variants, facilitate functional studies, and improve our understanding of their biological significance.

In this study, we developed a simple and efficient O_PCR_ based method, a flexible single-tube or dual-tube technique for cloning short or large gene/genomic fragments into any site within a plasmid. This protocol enables the insertion of single, dual, or multiple gene fragments into a suitable vector at any restriction site, regardless of fragment size. It can be performed with or without intermediate PCR purification steps. Furthermore, we introduced diverse site-directed mutations, including multiple substitution mutations at single and distinct loci, insertions, deletions, and a combination of these mutations in a single round of reaction. We also adapted our method to perform *de novo* synthesis of short gene fragments.

## RESULTS

Molecular cloning and mutagenesis are standard approaches for elucidating the functional relationships between genes and their encoding proteins. Given the indispensable role of these methods in molecular biology, there is a continuous need for improved strategies that are both efficient and user-friendly. In this study, we developed a simple and versatile PCR-based method for cloning, mutagenesis, and gene synthesis. The principle of the method relies on overlap extension PCR, which utilizes the 5’→3’ polymerase activity of DNA polymerase (Fig. 1). To simplify and validate our method, we first constructed a minimalistic plasmid, termed the mini cloning and expression plasmid (pMCE). The use of a small plasmid construct minimizes PCR-related errors and improves the transformation and transfection efficiencies. The pMCE, with the size of 3 kb, consists of essential regulatory elements, including the ori, the ampR gene, the T7 and CMV promoters, the bGH poly (A) signal sequence, and the MCS for cloning desired genes (Supplementary Fig. S1). Similar to other cloning plasmids, pMCE replicates in *E.coli* and generates a high copy number. It was used for validation of all our assays, including cloning, mutagenesis, and oligo assembly experiments.

**Figure 1.**
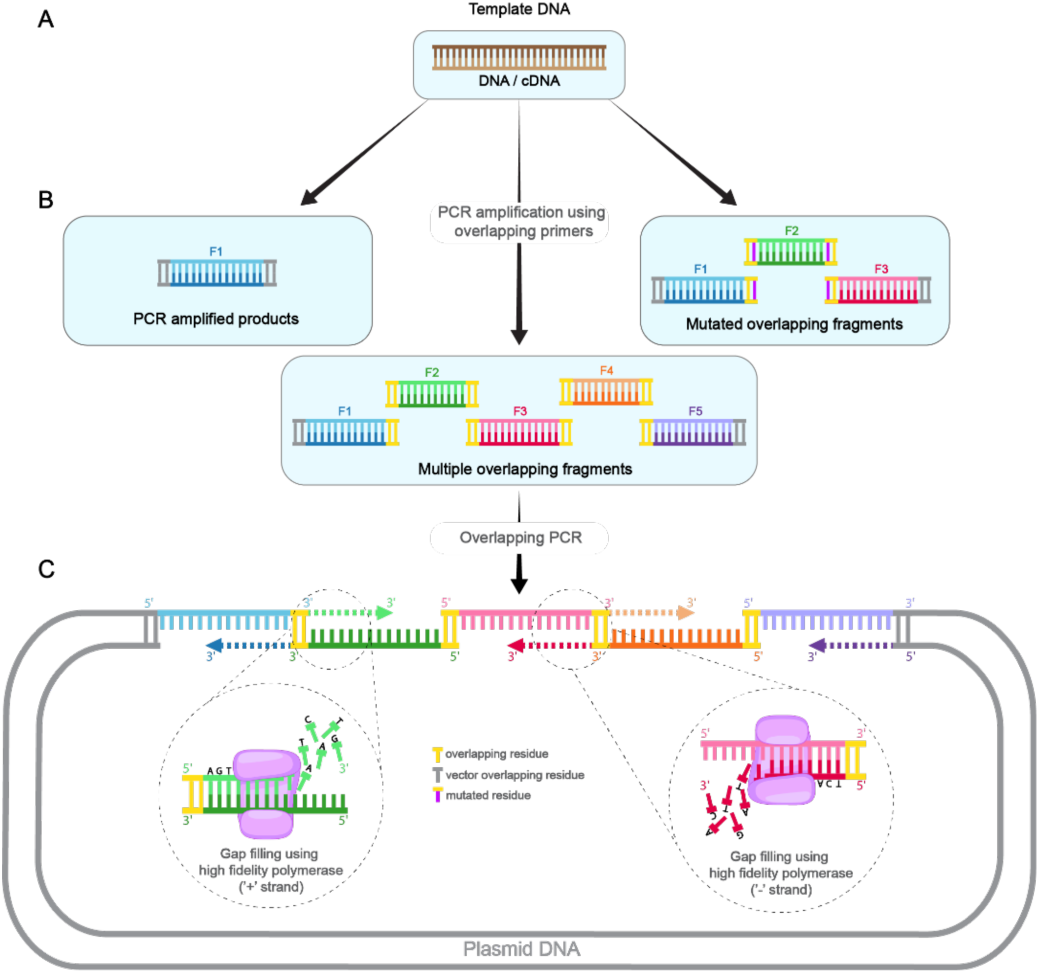
Schematic representation of the O_PCR_ based VSR method for cloning and site-directed mutagenesis. (**A**) DNA or cDNA serves as the initial template. (**B**) The desired gene/fragment is amplified using overlapping primers to generate overlapping sequences towards the 5’ and 3’ ends complementary to the plasmid DNA. The strategy enables the cloning of single or multiple gene fragments, including mutated products into plasmid DNA. In the case of multiple gene fragments, the first fragment contains a 5′ overlap with the plasmid, the last fragment contains a 3′ overlap with the plasmid, and the intermediate fragments carry 15 bp overlaps with their neighbouring fragments. (**C**) Overlap PCR reaction, the amplified gene fragments and linearized plasmid are combined with a high-fidelity polymerase, which extends the overlapping ends in the 5′ to 3′ direction, thereby filling the gaps and joining the adjacent fragments to generate a circular plasmid with a single nick at each junction.

We also prepared in-house competent cells using three distinct approaches: routine calcium chloride, ultracompetent cell, and electrocompetent cell preparation. Initially, the efficiency of the competent cells was analyzed by transforming them with varying concentrations of pMCE plasmid. All three methods yielded good transformation efficiency, but the ultracompetent and electrocompetent cells resulted in significantly higher colony numbers (data not shown).

### Efficient insertion of eGFP gene into pMCE validates O_PCR_ based cloning

We first cloned a 720 bp eGFP gene into the pMCE expression vector as a proof-of-concept. The pMCE vector was linearized using the blunt-end restriction enzyme EcoRV, and the resulting linearized plasmid was used for all validation experiments. The eGFP gene was PCR-amplified using overlapping cloning primers, resulting product containing 15 bp overlaps at both the 5’ and 3’ end of the plasmid. *In vitro* cloning was performed by PCR using 0.025 pmol of the linearized pMCE vector and either 0.025 pmol of the purified eGFP product or 0.5−1 µl of unpurified PCR product, along with a high-fidelity polymerase. Following the reaction, 1 µl of the PCR product was directly transformed into chemically competent *E. coli* DH5α cells without further purification. Colony PCR showed 7 out of 10 clones (70%) contained desired insert in the correct orientation, and no sequence errors were detected in the positive clones (Fig. 2). Further, transfection of these plasmids into HEK293T cells resulted in green fluorescence, confirming the successful insertion and expression of eGFP (Fig. 3L).

**Figure 2.**
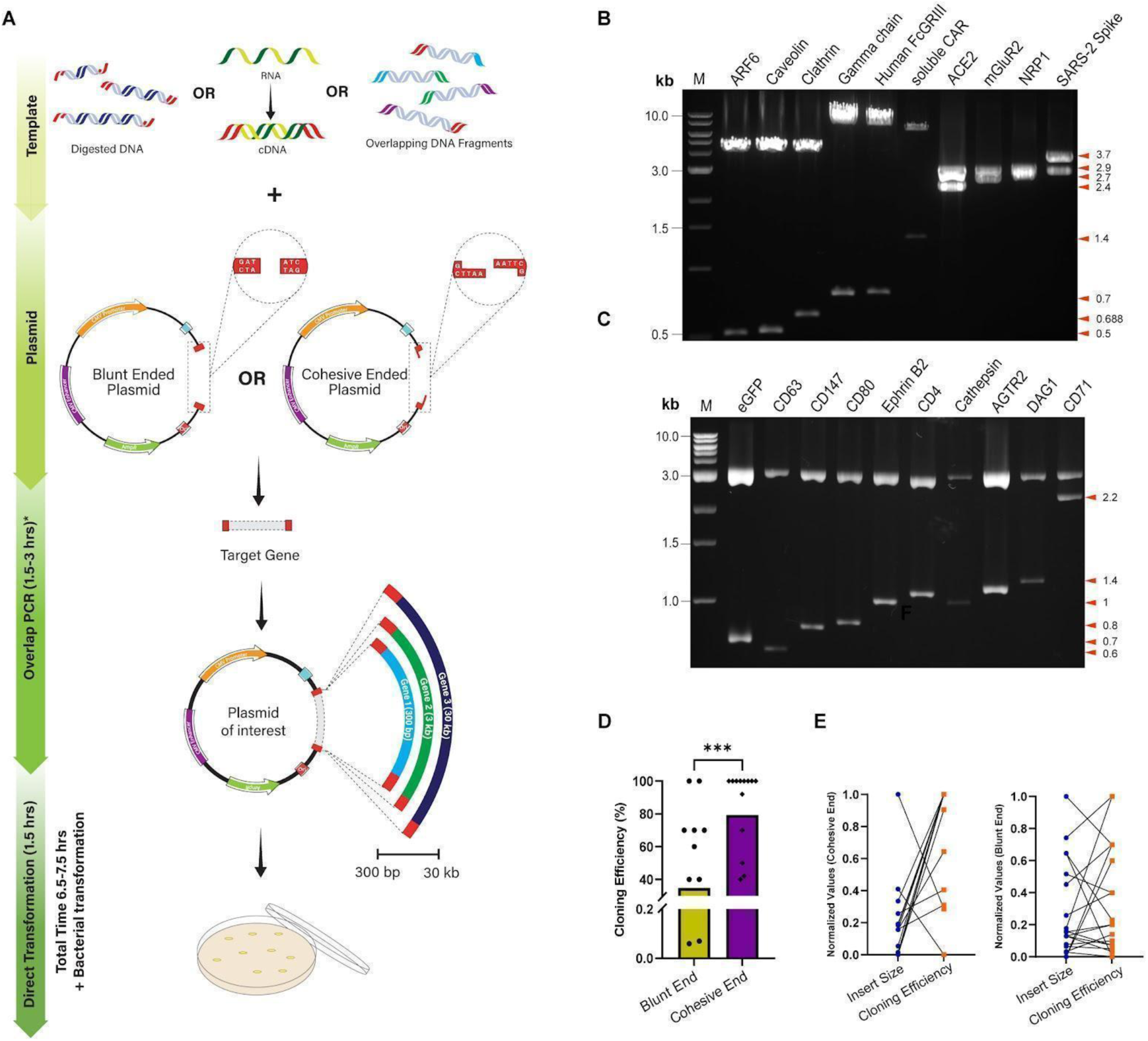
Ligase-free cloning of diverse genes using the VSR method (**A**) Schematic representation of gene cloning using O_PCR_, depicting the types of template DNA, plasmid vector, sizes of the gene, and total time employed in the method. Electrophoretic analysis of restriction digestion products loaded on 1% agarose gel (**B**) cloned using purified DNA products as templates (**C**) clones generated using un-purified PCR products as templates; Lane M–1 kb DNA molecular weight ladder. (**D**) Bar graph representing the comparison of cloning efficiency between blunt-end and cohesive-end digested plasmid DNA is plotted. (**E**) Comparison of insert size and cloning efficiency in cohesive-end and blunt-end cloning plotted as a slope graph.

**Figure 3.**
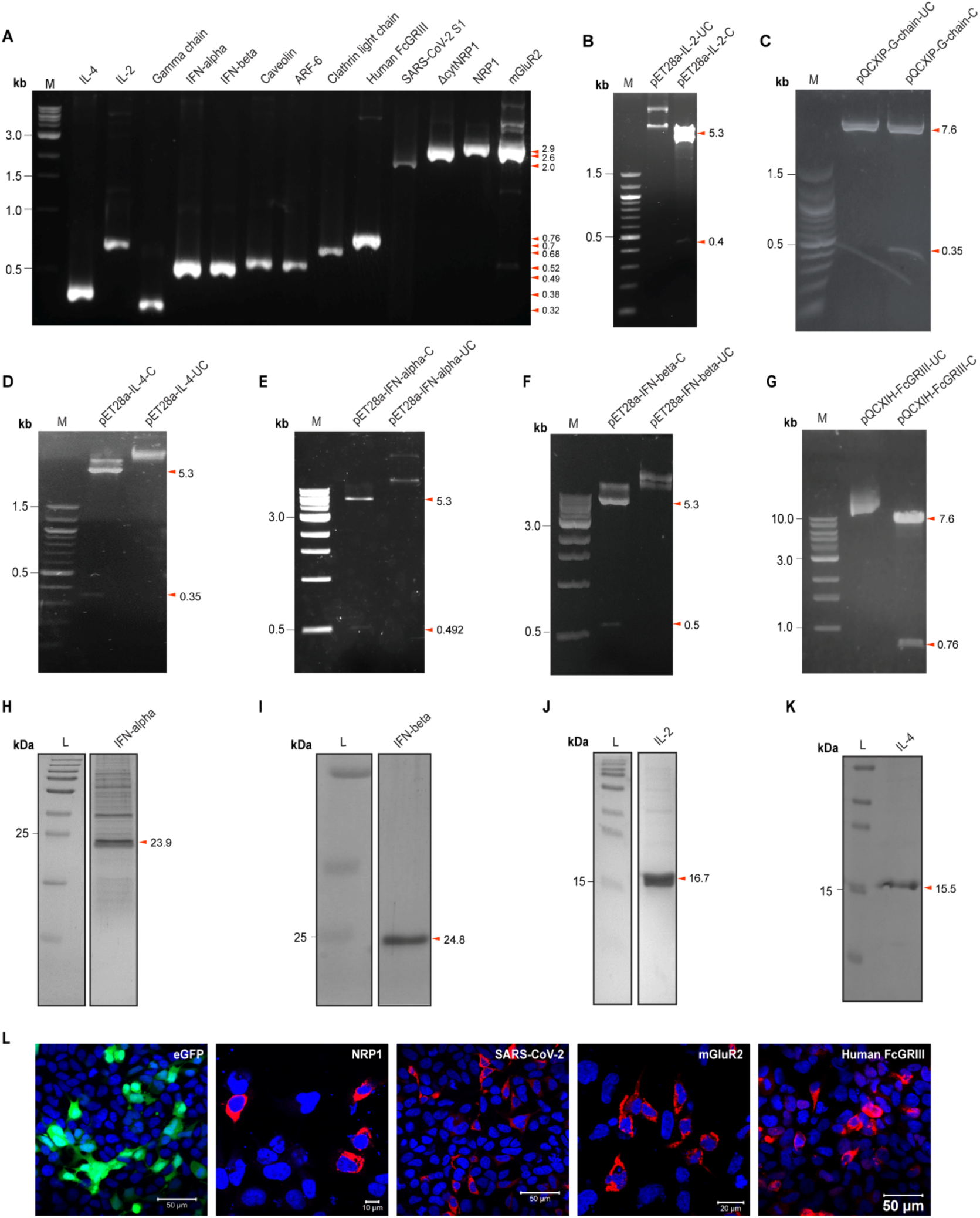
Cloning of genes using the VSR method. Agarose gel electrophoresis (1% agarose) of (**A**) PCR amplified product of 13 genes using specific overlapping primer sets. Restriction digestion analysis for the confirmation of the generated clones (**B**) IL2 (**C**) G-chain (**D**) IL4 (**E**) IFN-alpha (**F**) IFN-beta (**G**) FcGRIII; Lane M-1 kb DNA molecular weight marker. SDS-PAGE gel confirming protein expression of cloned genes (**H**) IFN-alpha (**I**) IFN-beta (**J**) IL-2 (**K**) IL4; Lane L-250 kDa protein size marker (**L**) Immunofluorescence staining of cells transiently expressing eGFP, NRP1, SARS-CoV-2 S1, mGluR2 and human FcGRIII using protein specific antibodies; scale bar= 10 um (NRP1), 20 um (mGluR2), 50 um (eGFP, SARS-CoV-2, Human FcGRIII).

### VSR is a versatile method for cloning of diverse genes of varying sizes

#### VSR method bypasses the challenges posed by traditional cloning methods

Next, the VSR protocol was applied to clone 35 distinct genes of varying fragment sizes (80 bp to 6 kb), including both viral and eukaryotic genes. These gene fragments were successfully inserted into desired vectors at various sites, using either blunt-end or cohesive-end cloning strategies involving single or double restriction digestion. The ease and flexibility of any cloning method largely depend on the availability of less stringent options for selecting a suitable plasmid vector, selection markers, and further downstream manipulations, along with the total time required to complete the protocol. To assess the adaptability of the VSR method, we used five different plasmid vectors of varying sizes: pMCE (3 kb), pTurboFP6335 (4.6 kb), pcDNA3.1+ (5.4 kb), pET28a (5.36 kb), and pQCXIP (7.2 kb). A key challenge in conventional cloning is the requirement for compatible restriction sites between the insert and the vector. However, the O_PCR_ used in VSR bypasses this need, as the insert does not require restriction digestion. Instead, the vector can be linearized using any restriction enzyme(s), blunt or cohesive, and either single or double digested, thereby significantly broadening the spectrum of suitable vectors. The linearized plasmid and PCR-amplified insert can be used directly for O_PCR_.

To simplify the method further, we used three distinct approaches to optimize gene cloning. In the first approach, PCR-amplified inserts were purified and then subjected to O_PCR_ using vectors digested with either blunt or cohesive end enzymes (Fig. 2A). The O_PCR_ products were directly transformed into ultra-competent *E. coli* cells without additional purification steps. Using this approach, we successfully cloned 17 diverse genes, including those encoding membrane-bound and secretory proteins, organelle markers, cytokines, and viral proteins, with fragment sizes ranging from 80 bp to 3 kb (Fig. 2B,3; Supplementary Table S1). Among them, 10 genes were cloned into vectors digested using cohesive ends generated by double digestion, 6 genes into blunt-end digested vectors using single restriction enzyme, and 1 gene using a single cohesive-end cutter. Remarkably, positive clones were obtained for all genes, with cloning efficiencies ranging from 70% to 100% (Fig. 2), and all successful clones were recovered in the first attempt. Moreover, vectors digested with cohesive-end restriction enzymes showed higher cloning efficiencies compared to those digested with blunt-end enzymes (Fig. 2D). This reduced efficiency in blunt-end cloning likely resulted from direct purification using PCR cleanup kits rather than gel extraction, which resulted in residual undigested or partially digested plasmid DNA. However, cloning with gel-purified, restriction-digested plasmids significantly improved efficiency by effectively removing incompletely digested or undigested vectors, thereby reducing background colonies and enhancing overall cloning efficiency.

A common bottleneck in cloning workflows is the low yield of purified inserts following PCR cleanup or gel extraction, which often compromises ligation efficiency. This challenge becomes more pronounced when insert bands are faint, as purification of such low-abundance products can result in significant material loss, ultimately reducing cloning efficiency or yielding no clones. Additionally, DNA elution from agarose gels may pose a potential risk of ethidium bromide-induced mutations (60). To circumvent this problem, we used our second approach, we attempted to clone 16 distinct genes (300 bp to 3 kb): 4 from DNA templates (previously cloned plasmid DNA) and 12 genes directly amplified from a cDNA library obtained from different mammalian cell lines. Interestingly, we obtained positive clones for all 16 genes in the first attempt. Although the efficiency for cDNA-derived clones was less, the genomic DNA-derived clones yielded an efficiency of 60% to 100% (Fig. 2C). Despite lower efficiencies in some cases, our results demonstrate that this streamlined protocol enables effective cloning from limited or low-quality DNA sources. Furthermore, cloning efficiency appeared unaffected by insert size for both blunt and cohesive-end cloning across a range of 80 bp to 3 kb (Fig. 2E).

#### VSR method enables the cloning of multiple gene fragments in a single reaction

In our third approach, to introduce shorter regions like tags and homologous arms, the desired fragments were amplified as two or more amplicons with overlapping regions and stitched together into a single DNA fragment using O_PCR_. For instance, we stitched two short fragments into the pMCE vector; the first fragment contained a 270 bp of domain 11 of eGFP (eGFP11) along with a glycine linker, and the second fragment contained 711 bp of the Fc domain of human IgG (Fig. 4A). These two gene fragments were individually amplified using overlapping cloning primers (Fig. 4B,C) and the amplified fragments were stitched into a single fragment using O_PCR_ (Fig. 4D) and then cloned into pMCE vector. The positive clones were confirmed using restriction digestion, which yielded the expected size of ∼984 bp (Fig. 4E). Upon transiently expressing on HEK293T cells, we were able to visualize a positive green fluorescence signal by immunofluorescence staining using an antibody against the human IgG-Fc tag (Fig. 4F). Additionally, immunoblot analysis showed a band at ∼29 kDa, consistent with the predicted molecular weight of the fusion protein (Fig. 4G). One of the challenges in cloning of certain vectors used for homologous recombination in bacterial system is the requirement of homologous arms flanking the desired gene. Traditionally, left and right arm are cloned into a vector system with restriction site to support clone the desired gene. However, this approach is time-consuming and not ideal, as it requires introduction of additional mutation to create restriction site between the homologous arms. To avoid these mutations, here, we amplified a 1 kb kanamycin gene and 200 bp left and right arms as overlapping products and stitched them using the VSR method (Fig. 4H-M). We successfully cloned the kanamycin gene in between the homologous arms, and the plasmid conferred resistance against kanamycin. This data clearly shows that the VSR method can facilitate efficient assembly and cloning of multiple sized fragments without the addition of any restriction site into a gene or vector.

**Figure 4.**
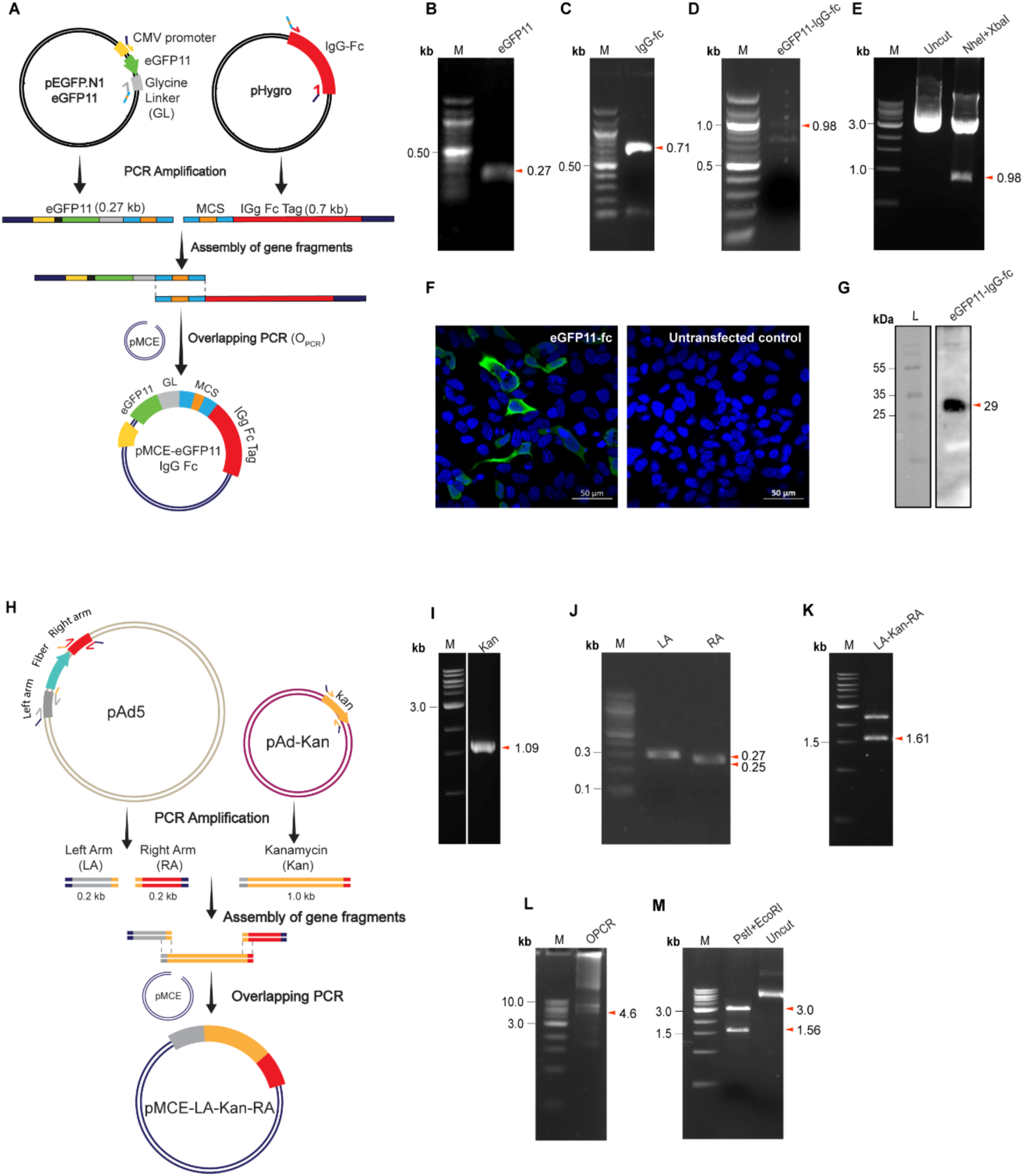
Introduction of a short linker fragment into a gene using VSR method (**A**) Schematic representation of cloning the eGFP11 gene fragment with the addition of multiple cloning sites (MCS), glycine linker (GL), and IgG-Fc tag into the pMCE expression vector. Agarose gel electrophoresis (1% agarose) of PCR-amplified product of (**B**) eGFP11, (**C**) IgG-Fc, (**D**) eGFP11-IgG-Fc, using overlapping primer sets; Lane M–100 bp DNA molecular weight marker (**E**) Restriction digestion analysis of eGFP11-IgG-Fc; Lane M-1 kb DNA molecular weight marker (**F**) Fluorescence imaging of cells transiently expressing eGFP11-IgG-Fc compared to the un-transfected control; scale bar = 50 µm (**G**) Western blot analysis confirming the protein expression of eGFP11-IgG-Fc; Lane L–250 kDa protein size marker. (**H**) Schematic representation of cloning kanamycin resistance gene (kan) by addition of 200 bp homologous arms upstream (left arm, LA) and downstream (right arm, RA) into pMCE expression vector. Agarose gel electrophoresis (1% agarose) of PCR amplified product of (**I**) Kanamycin; Lane M-1 kb DNA molecular weight marker (**J**) Left Arm (LA) and Right Arm (RA); Lane M-100 bp DNA molecular weight marker (**K**) LA-Kan-RA fragment; Lane M-1 kb DNA molecular weight marker (**L**) overlap PCR product of pMCE-LA-Kan-RA; Lane M-1 kb DNA molecular weight marker (**M**) Restriction digestion product of pMCE-LA-Kan-RA positive clone; Lane M-1 kb DNA molecular weight marker.

Next, we applied our method for assembling more than three gene fragments of varying sizes using VSV minigenomes as a proof-of-concept. First, for constructing the VSV-Nucleoprotein-Phosphoprotein minigenome (VSV-NP), we PCR amplified full-length (2.25 kb) of the VSV-NP gene, 140 bp of the VSV-trailer sequence, and 150 bp of the hepatitis delta virus-ribozyme (HDVr) independently, and all these three fragments were cloned into the pMCE vector in a single reaction (Fig. 5A-D). Similarly, the large polymerase gene of VSV (VSV-L) of full-length (6.4 kb) was amplified as three fragments of 2.8 kb, 2 kb, and 1.6 kb, which were assembled along with the VSV-leader sequence (110 bp) and HDVr (150 bp) into the pMCE vector (Fig. 5F-H). Interestingly, positive clones were obtained in the first attempt, and the resulting positive clones were confirmed using dual and triple cut restriction digestion. In both the VSV-minigenomes, the desired restriction digestion pattern was observed (Fig. 5E, I), confirming the correct assembly and integrity of the plasmid construct.

**Figure 5.**
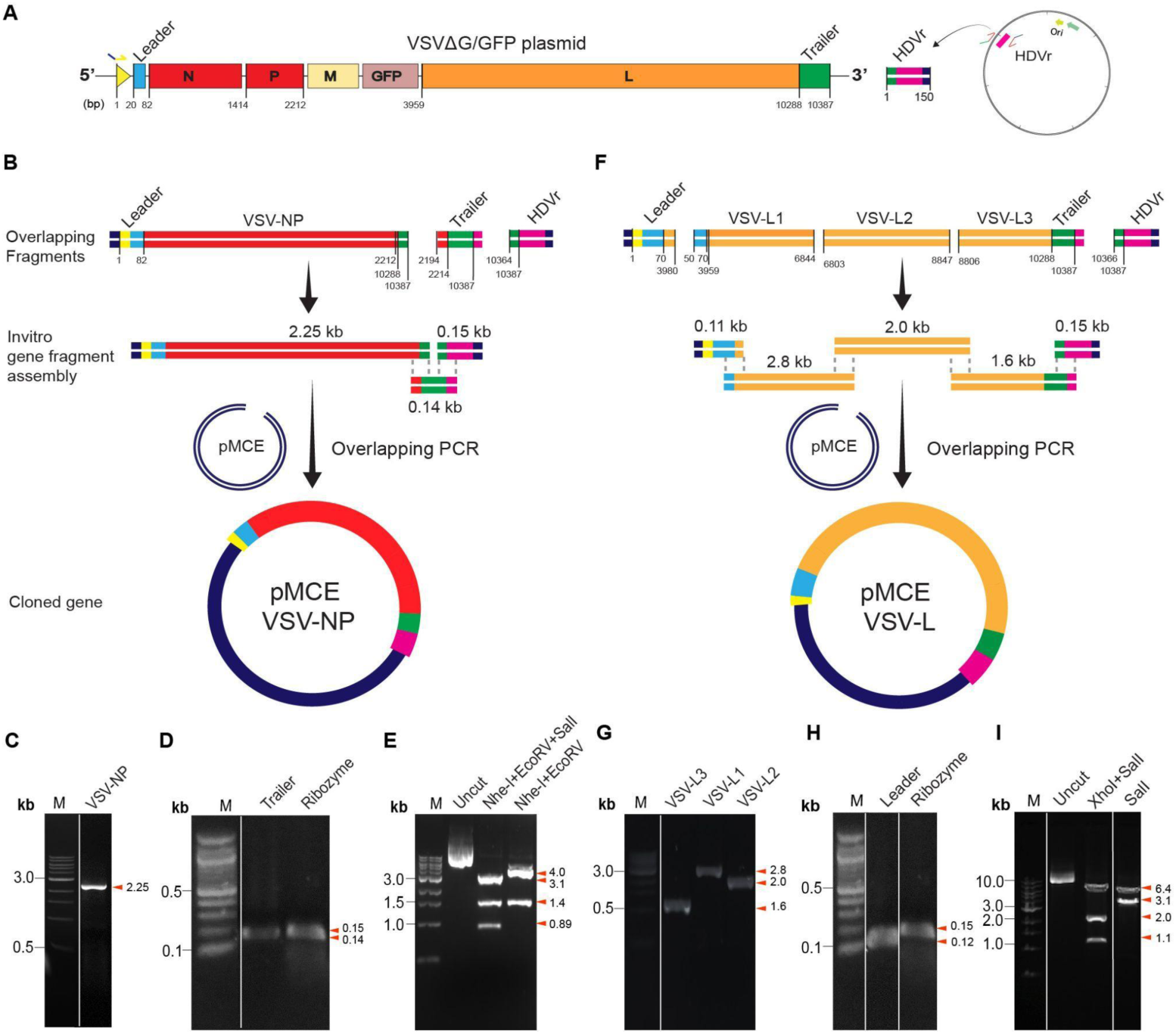
Stitching and cloning of more than three fragments of varying sizes in a single tube and generation of minigenome plasmids of VSV. Schematic representation of (**A**) plasmid maps of VSVΔG/GFP and hepatitis delta virus ribozyme (HDVr). (**B**) Cloning of vesicular stomatitis virus (VSV) minigenome plasmids-VSV-Nucleoprotein-Phosphoprotein (VSV-NP). Agarose gel electrophoresis (1% agarose) of PCR amplified product of (**C**) VSV-NP gene; Lane M–1 kb DNA molecular weight marker (**D**) trailer and HDVr; Lane M-100 bp DNA molecular weight marker. (**E**) Restriction digestion mapping of pMCE-VSV-NP; Lane M–1 kb DNA molecular weight marker. (**F**) Schematic representation of cloning the VSV-Large Polymerase protein (VSV-L) into the pMCE expression vector. Agarose gel electrophoresis (1%) of PCR amplified product of (**G**) VSV-L1, VSV-L2, VS-L3; Lane M-1 kb DNA molecular weight marker (**H**) Leader and HDVr gene fragments; M–100 bp DNA molecular weight marker. (**I**) Restriction digestion analysis of pMCE-VSV-L plasmid; Lane M–1 kb DNA molecular weight marker. Distinct lanes cropped from one gel image have been assembled as a composite image and separated by white lines and correspond to the marker run on the same gel.

#### Cloning of Large and Difficult-to-Amplify Viral Genomes Using the VSR Method

Cloning large genes (>7 kb) or genomes that are difficult to amplify present a significant challenge using conventional cloning methods. To demonstrate the robustness of the VSR method, we successfully cloned the full-length genome (∼10.3 kb) of vesicular stomatitis virus lacking the glycoprotein gene but expressing GFP (VSV-ΔG/GFP). As VSV is a negative-sense RNA virus, the viral RNA was first isolated and reverse-transcribed into cDNA. The full-length genome was then amplified as five overlapping fragments ranging from 1.0 kb to 2.9 kb and a 150 bp of HDVr sequence (Fig. 6A-C). Then, 0.05 pmol of each of the six gene fragments, along with the pMCE vector backbone, were subjected to O_PCR_ (Fig. 6D). The final assembled mixture was directly transformed into *E. coli* DH10B using electroporation. We obtained positive clone in the first attempt, which were further confirmed by restriction digestion analysis (Fig. 6E).

**Figure 6.**
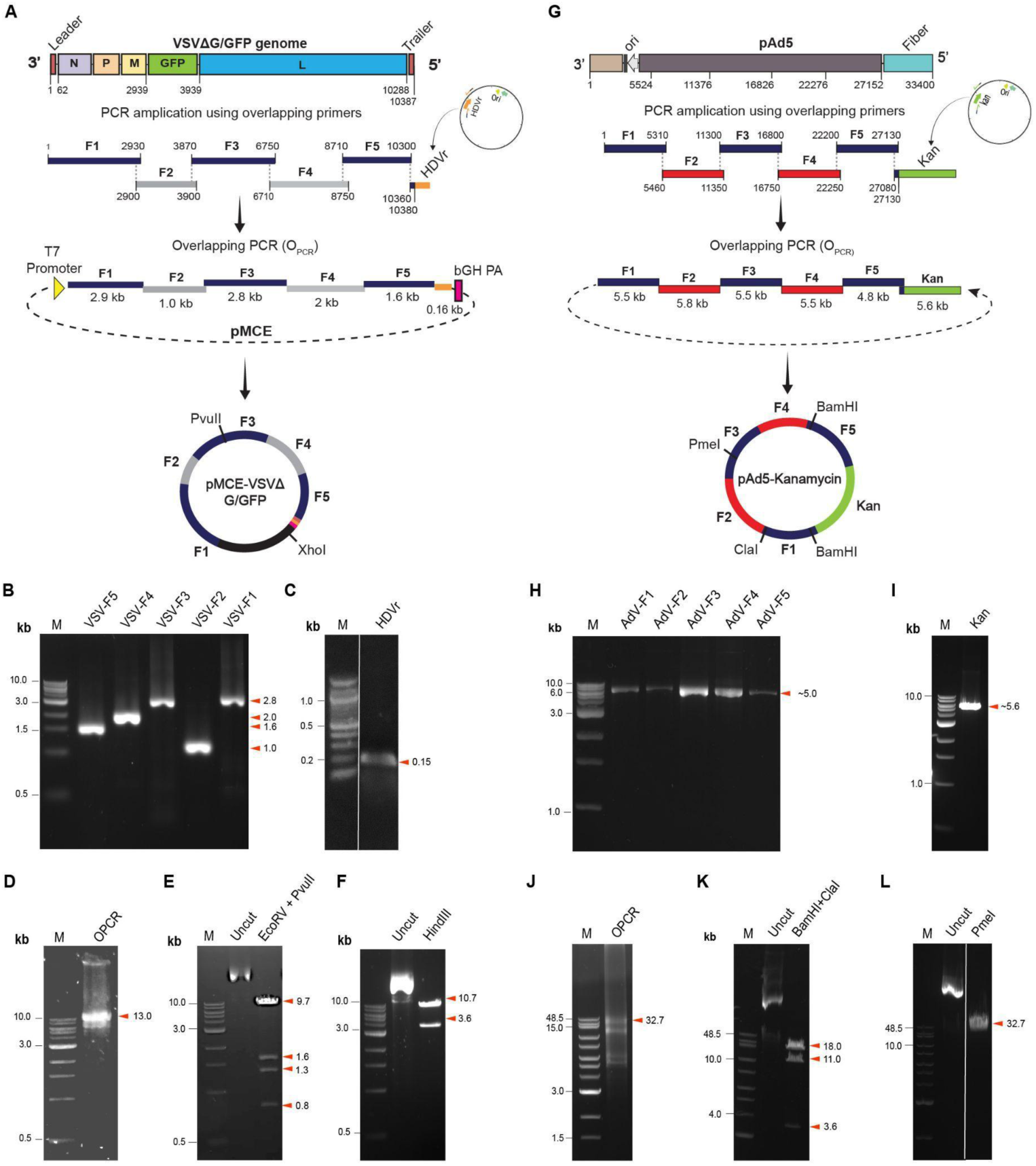
Cloning of large genomic fragments using O_PCR_ (**A**) Schematic representation of cloning a 10.3 kb sized VSVΔG/GFP genome into a pMCE expression vector. Electrophoretic analysis of PCR-amplified products (**B**) VSV fragments–VSV-F5, VSV-F4, VSV-F3, VSV-F2, and VSV-F1 loaded on 1% agarose gel; M–1 kb DNA molecular weight marker; (**C**) hepatitis delta virus ribozyme (HDVr) loaded on 1.5% agarose gel; M–100 bp DNA molecular weight marker **(D**) O_PCR_ product mix of VSV fragments (F1-F5), HDVr and pMCE loaded on 1% agarose gel; M–1 kb DNA molecular weight marker (**E,F**) restriction digestion mapping of the plasmid DNA isolated from the positive clone loaded on 1% agarose gel; M–1 kb DNA molecular weight marker. (**G**) Schematic representation of cloning a 32.7 kb sized Ad5 genome by replacing the fiber gene with a kanamycin resistance gene. Electrophoretic analysis of PCR-amplified products (**H**) Adenovirus fragments–AdV-F1, AdV-F2, AdV-F3, AdV-F4, and AdV-F5 (**I**) Kanamycin resistance gene loaded on 1% agarose gel; M–1 kb DNA molecular weight marker (**J**) O_PCR_ product mix of AdV fragments (F1-F5) and kanamycin gene; restriction mapping analysis of plasmid DNA isolated from the positive clone using (**K**) dual cutter and (**L**) unique cutter enzyme; loaded on 0.8% agarose gel; M–1 kb extended DNA molecular weight ladder. Distinct lanes cropped from a single gel image have been assembled as a composite image. In such grouped images, the lanes are separated by ‘white’ dividing lines and correspond to the marker run on the same gel.

To further evaluate the capacity of this method for cloning even larger genomes, we attempted to clone a 33 kb genome of human adenovirus type 5 (Ad5), a double-stranded DNA virus (Fig. 6G). The pAd plasmid carrying the Ad5 genome was amplified into five overlapping fragments, each ranging from 4.8 kb to 5.8 kb, with 100 bp of overlap between the adjacent fragments (Fig. 6H). To show the efficiency of the cloning technique, the fiber gene (1.8 kb) of the adenovirus was replaced with kanamycin resistance gene (1 kb). The kanamycin cassette was prepared with overlapping regions corresponding to the left and right arms of the fiber gene (Fig. 6I). The five adenoviral fragment and the kanamycin cassette was assembled using O_PCR_ (Fig. 6J). One microliter of the assembled product was electroporated into *E. coli* DH10B bacteria. Interestingly, one out of six clones screened exhibited the correct size and orientation in the first attempt. Restriction mapping analysis showed the expected 32.7 kb band, which confirmed the successful assembly of adenovirus gene fragments as well as the replacement of the fiber gene with the kanamycin cassette (Fig. 6K,L). Upon transforming the plasmid into *E coli* DH10B bacteria, it conferred resistance against kanamycin. Furthermore, the deletion of the fiber gene was confirmed by transfecting the plasmid in HEK293T cells which showed no expression of fiber protein.

### VSR is a Versatile Platform for Introducing Diverse Mutations into a Gene Without the Need for Prior Cloning

#### Validation of the VSR Method Using a Single Substitution Mutation

Site-directed mutagenesis is a powerful tool for investigating gene function by enabling the precise introduction of nucleotide changes. It is also widely applied in the engineering of recombinant proteins and organisms to enhance or modify function. To demonstrate the efficiency of the VSR method for SDM, we introduced targeted mutations into the S1 domain of the SARS-CoV-2 spike gene, as well as the human dipeptidyl peptidase 4 (hDPP4) and human angiotensin-converting enzyme 2 (hACE2) genes as proof-of-concept targets. As an initial step to evaluate and optimize the method, a single substitution mutation (H403S) was introduced into the spike gene. The target region was amplified in two overlapping fragments (FR1 and FR2) using overlapping mutagenic primers (as shown in Fig.7). These fragments were then assembled and cloned into the pMCE plasmid backbone by O_PCR_. One microliter of the O_PCR_ product was directly transformed into *E. coli* DH5α competent cells (Fig. 8A). Interestingly, all 25 screened clones (100%) carried the desired mutation, with no off-target changes detected by Sanger sequencing (Fig. 8Bi).

**Figure 7.**
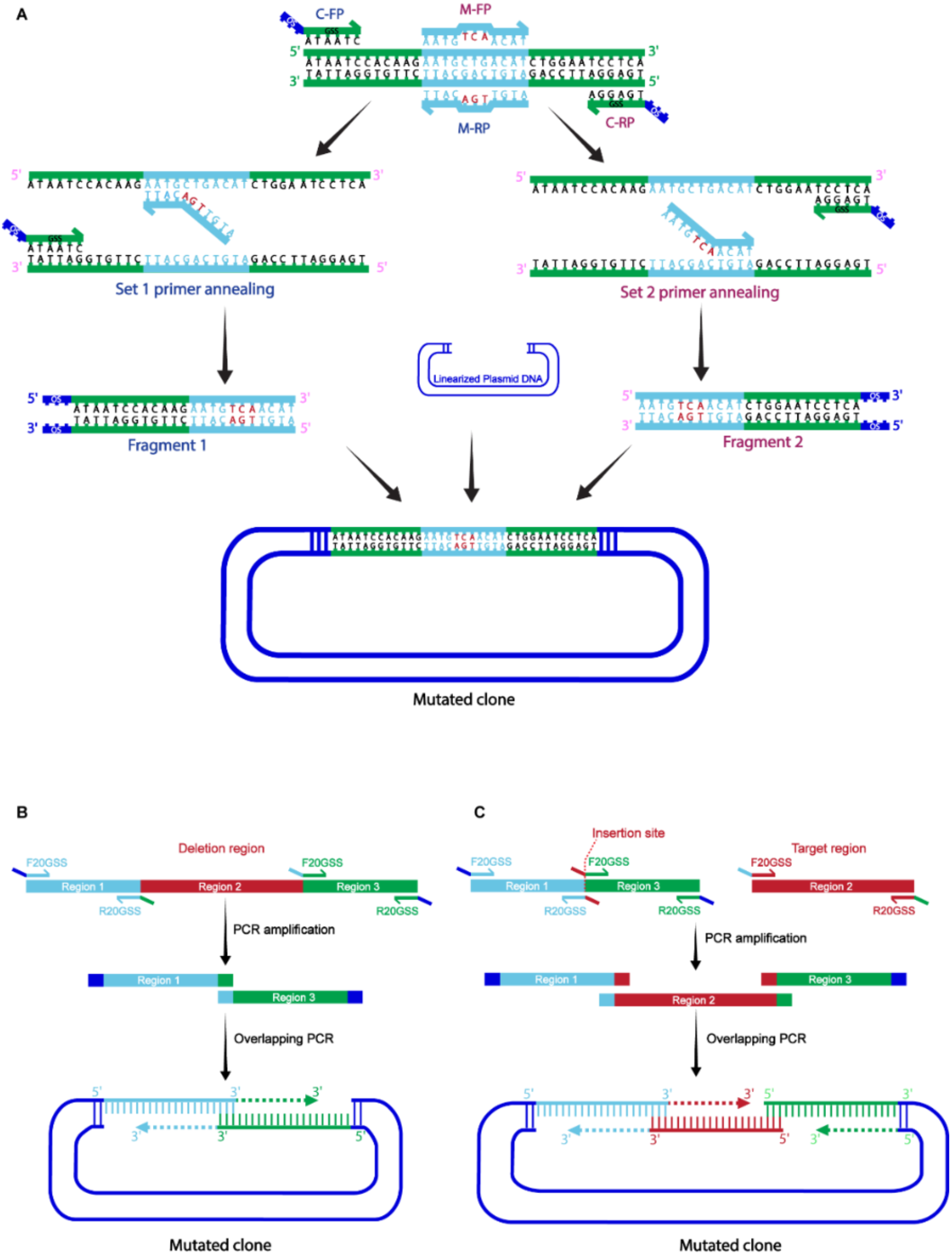
(**A**) Schematic representation of primers designed for introducing mutations. C-FP and C-RP-forward and reverse primers for cloning; M-FP and M-RP-forward and reverse primers for introducing mutations. Set-1 primers consist of C-FP and M-RP indicated in blue colour and Set-2 primers consist of C-FP and M-RP indicated in pink colour. (**B**) Schematic representation for introducing large insertion and deletion mutation. F-Forward primer, GSS-Gene Specific Sequence. The target region to be deleted or inserted is shown in red colour.

**Figure 8.**
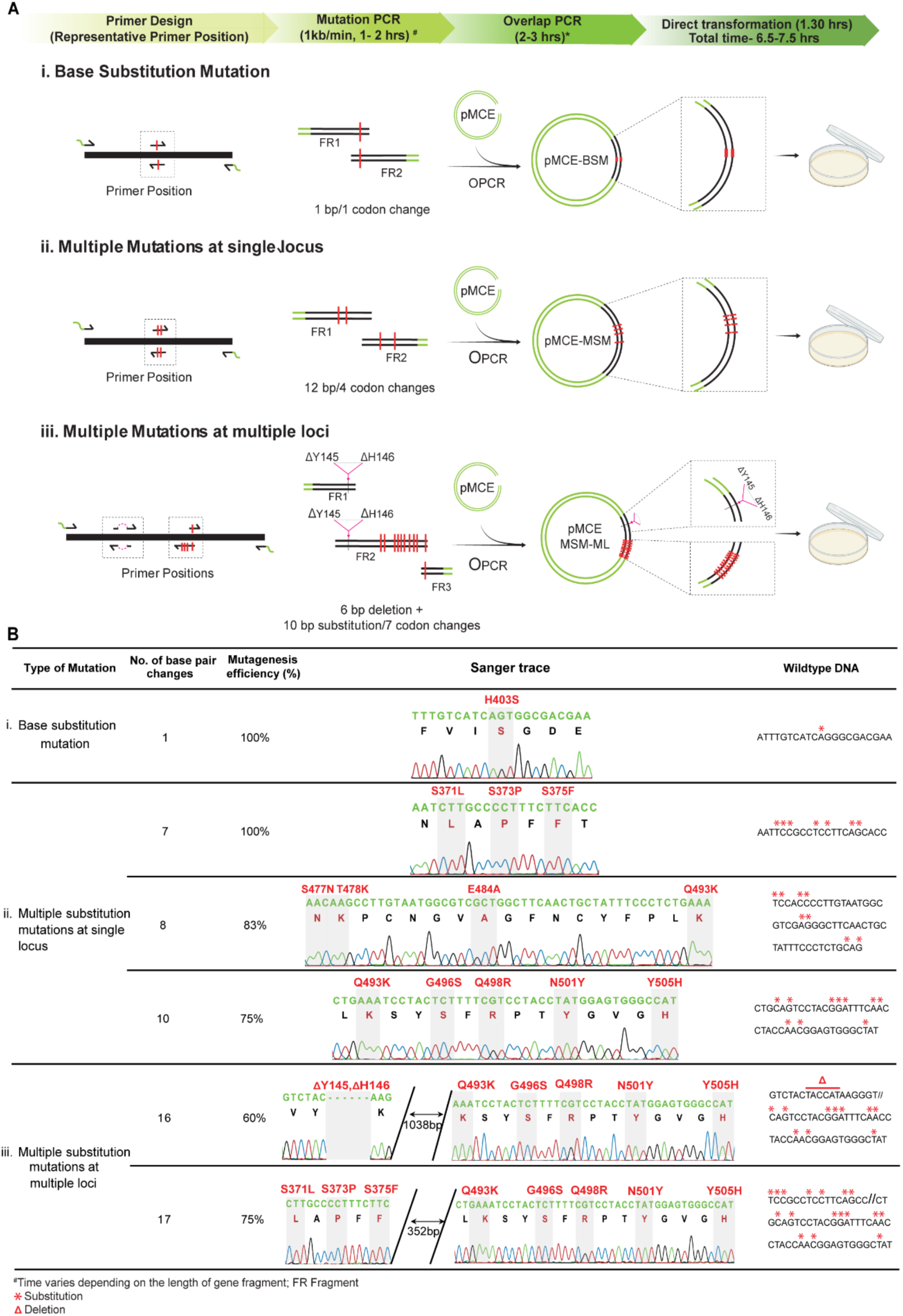
Incorporation of single and multiple substitution mutations using O_PCR_ (**A**) Schematic representation of introduction of single substitution mutation and multiple substitution mutations at single and multiple loci. (**B**) Tabulation depicting the detailed information of the number of mutations, mutagenesis efficiency, and Sanger trace analysis in comparison with wild-type DNA. Nucleotides marked with red asterisk marks represent mutated nucleotides.

#### VSR Method Allows Multiple Nucleotide Substitutions at a Single Locus

While several protocols exist for introducing single-point mutations, only a limited number support the incorporation of multiple nucleotide changes in a single reaction [76]. To evaluate the versatility of our method, we next introduced multiple substitutions at a single locus. Specifically, we generated three distinct substitution mutations involving 7 nt (corresponding to 3 amino acids), 8 nt (4 amino acids), and 10 nt (5 amino acids), each introduced independently in a single reaction (Fig. 8Bii). Interestingly, sequencing analysis revealed that the plasmids containing the 7 nt substitution exhibited 100% mutagenesis efficiency. Constructs with 8 nt and 10 nt substitutions showed efficiencies of 83% and 75%, respectively. These results highlight the reliability of the VSR method in accommodating multi-nucleotide substitutions with high accuracy and efficiency.

#### VSR Method Allows the Simultaneous Introduction of Mutations at Distinct Loci

We further extended the assay to introduce multiple mutations, both substitutions and deletions, at distinct loci within a gene. Two sets of experiments were performed. In the first, we simultaneously introduced an 8 nt substitution at one site and a 10 nt substitution at a second site located approximately 300 bp apart. In the second experiment, a 10 nt substitution was combined with a 6 bp deletion at two sites separated by 1050 bp. Remarkably, positive clones were obtained on the first attempt in both cases, with mutagenesis efficiencies of 60% and 75%, respectively (Fig. 8Biii). These results highlight the efficiency and flexibility of the VSR method in enabling the introduction of complex, multi-site mutations in a single reaction.

To further assess the versatility of our approach for mutating both terminal regions and multiple internal sites across a gene, we applied the VSR method to two full-length human genes: hDPP4 (2.3 kb) and hACE2 (2.4 kb). First, to introduce a single mutation at the 3′ end of the hDPP4 gene (Fig. 9A), we used overlapping mutagenic primers to amplify two fragments: a 2.22 kb fragment and a short 80 bp fragment (Fig. 9B, C). These fragments were successfully assembled and cloned into the pMCE vector via overlap PCR (O_PCR_). Sanger sequencing confirmed 100% mutagenesis efficiency with no off-target nucleotide changes (Fig. 9D). Next, we introduced six nucleotide substitutions at the 3′ end of hDPP4 using a similar strategy. The gene was amplified into three overlapping fragments (F1: 2.1 kb; F2: 101 bp; F3: 131 bp) (Fig. 9E-G), and sequencing analysis confirmed the accurate incorporation of all six mutations without any errors (Fig. 9H).

**Figure 9.**
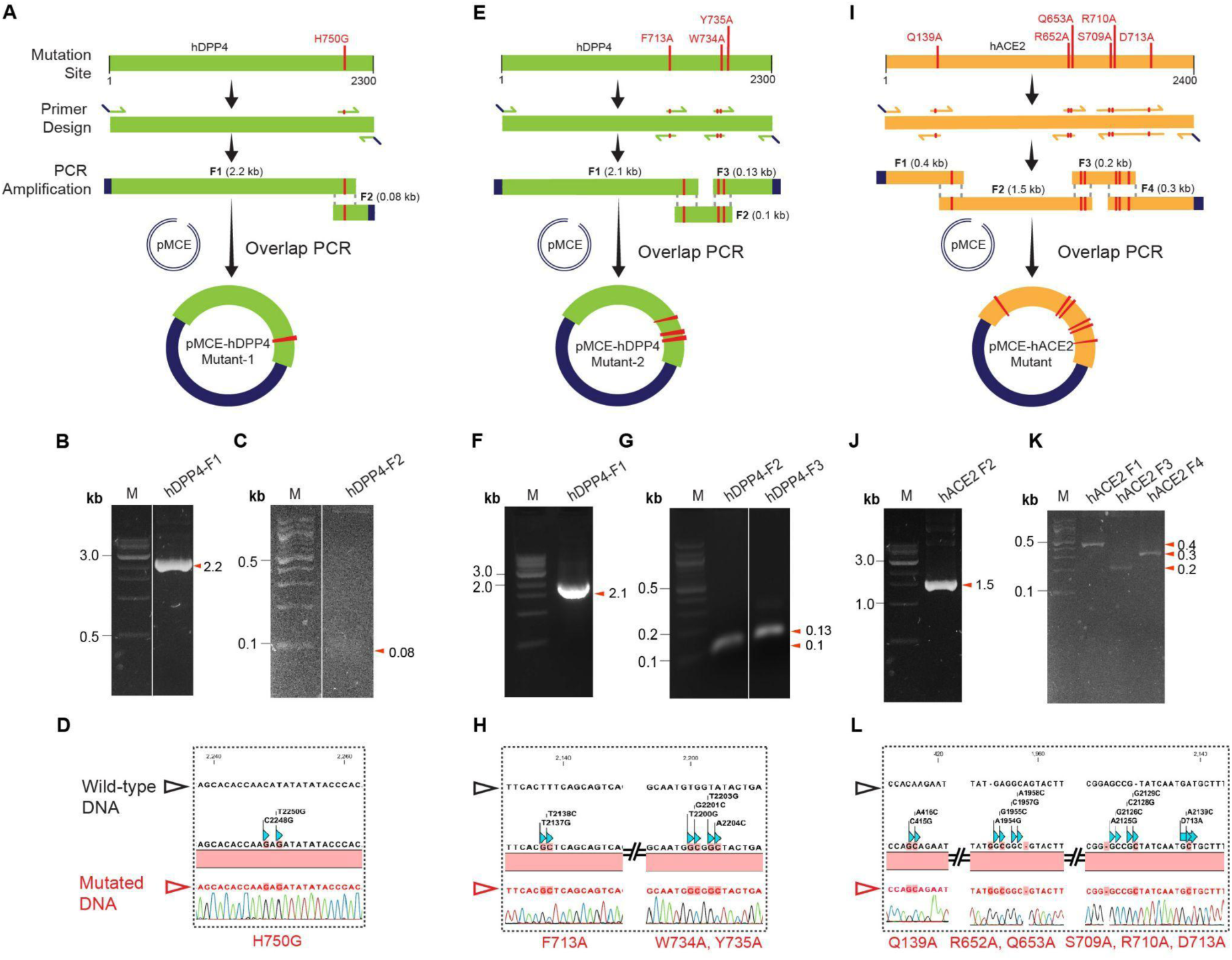
Introduction of multiple substitution mutations in the hDPP4 and hACE2 genes. (**A**) Schematic representation of introducing H750E substitution mutation in the human DPP4 receptor (mutant-1). Agarose gel electrophoresis of PCR amplified products of (**B**) hDPP4 F-1; Lane M–1 kb DNA molecular weight marker (**C**) hDPP4 F-2; Lane M–100 bp DNA molecular weight marker. (**D**) Sanger trace analysis of pMCE-hDPP4-Mutant-1 with reference to hDPP4 WT. (**E**) Schematic representation of introducing F713A, W734A, and Y735A mutations in the human DPP4 receptor (mutant-2). Agarose gel electrophoresis of PCR amplified products of (**F**) hDPP4 F-1; Lane M–1 kb DNA molecular weight marker (**G**) hDPP4 F-2, F-3; Lane M–100 bp DNA molecular weight marker. (**H**) Sanger trace analysis of pMCE-hDPP4-Mutant-2 with reference to hDPP4 WT. (**I**) Schematic representation of introducing Q139A, R652A, Q653A, S709A, R710A, and D713A mutations in the human ACE-2 receptor. Agarose gel electrophoresis of PCR amplified products of (**J**) hDPP4 F-1 (**K**) hDPP4 F-2, F-3; Lane M–1 kb DNA molecular weight marker. (**L**) Sanger trace analysis of pMCE-hACE-2-Mutant with reference to hACE-2 WT. Distinct lanes cropped from one gel image have been assembled as a composite image and separated by white lines and correspond to the marker run on the same gel.

We then extended this strategy to introduce 11 nt substitutions at three distinct loci within the hACE2 gene (Fig. 9I). Using overlapping mutagenesis primers, the gene was amplified into four overlapping fragments (F1: 0.4 kb; F2: 1.5 kb; F3: 0.2 kb; F4: 0.2 kb), which were assembled and cloned into the pMCE vector (Fig. 9J, K). Sequencing analysis revealed 100% efficiency, with all intended mutations correctly incorporated and no unintended nucleotide changes detected (Fig. 9L). Together, these results demonstrate the high efficiency and precision of the VSR method in introducing multiple mutations at multiple loci across an entire gene. This approach effectively overcomes a major limitation of conventional SDM methods, which often struggle to introduce multiple, spatially separated mutations in a single round of mutagenesis [61,76].

#### Efficient Insertion and Deletion Mutagenesis Using the VSR Method

Finally, we tested the ability of the method to introduce insertion and deletion mutations. Similar to the previous experiments, the desired short insertions (H496) and deletions (ΔY145, ΔH146) were incorporated into the primers, and gene fragments were PCR amplified independently using overlapping mutagenic primers (Fig. 7B,C). The amplified fragments were cloned into a pMCE vector using O_PCR_ and the product was directly transformed into *E. coli* (Fig. 10A). Sanger sequencing of 30 clones confirmed the presence of the desired insertion in all cases (100% efficiency), with no nonspecific mutations detected (Fig. 10Bi). Similarly, for the deletion mutation, all the 5 clones screened showed the correct mutation with no errors observed (Fig. 10Bii).

**Figure 10.**
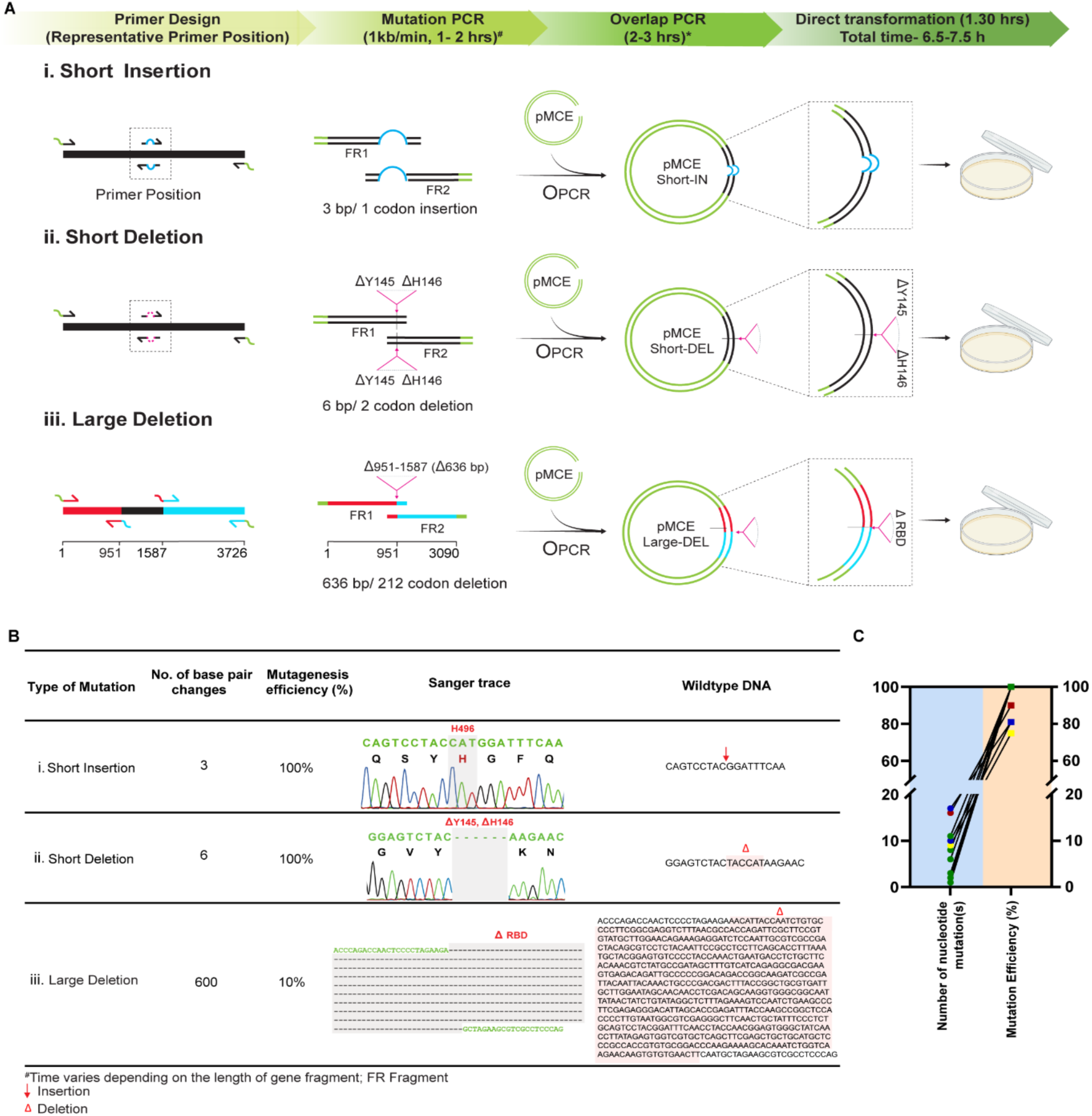
Incorporation of insertion and deletion mutations using O_PCR_. (**A**) Schematic representation of introduction of short insertion, short and large deletion mutations. (**B**) Tabulation depicting the detailed information of number of mutations, mutagenesis efficiency and Sanger trace analysis in comparison with the wild type DNA. Nucleotides highlighted in red colour are mutated. (**C**) Correlation of mutagenesis efficiency with the number of mutations incorporated using O_PCR._ Each coloured circle and the square in the graph represent the number of mutations incorporated corresponding to the mutagenesis efficiency.

To generate a large deletion mutation, two independent PCR reactions were performed using primer sets as illustrated in Fig. 7C, 10Aiii. To exclude the targeted region, the forward primer of the first set was designed with an overlapping cloning primer (20GSS), while the corresponding reverse primer contained 15 nucleotides immediately downstream of the deletion site. In parallel, the forward primer of the second set included 15 nucleotides immediately upstream of the deletion site, and the reverse primer was the overlapping cloning primer (20GSS). These two fragments were amplified separately and then joined into a single, continuous DNA fragment which lacks the desired deletion region. The overlap between the 3′ end of the first fragment and the 5′ end of the second fragment enabled smooth joining, effective removal of the targeted sequence as well as simultaneous cloning by O_PCR_ (Fig. 10Aiii). Positive clones were confirmed by Sanger sequencing, and the desired deletion construct was successfully obtained on the first attempt (Fig. 10Biii).

#### De novo gene synthesis using the VSR method

In addition to gene cloning and SDM, we evaluated the potential of the VSR method for synthesis of gene fragments using overlapping oligonucleotides (Fig. 11A). To assess this, we initially assembled oligonucleotides of 50 bp size in progressively increasing numbers and obtained amplicons of 94 bp, 160 bp, 311 bp, 455 bp, 603 bp and 702 bp. This showed that our approach can be efficiently used for synthesising a wide range of shorter fragments (Fig. 11B). To further determine the adaptability of the method for synthesizing gene fragments and subsequent cloning, we synthesized a 600 bp gene fragment encoding the receptor-binding domain (RBD) of the SARS-CoV-2 omicron variant using 18 overlapping oligonucleotides, each with a 15-22 bp overlap (Fig. 11C,D). The synthesized fragment was cloned into pMCE vector, and all 10 sequenced clones were accurate, demonstrating 100% cloning efficiency (Fig. 11E).

**Figure 11.**
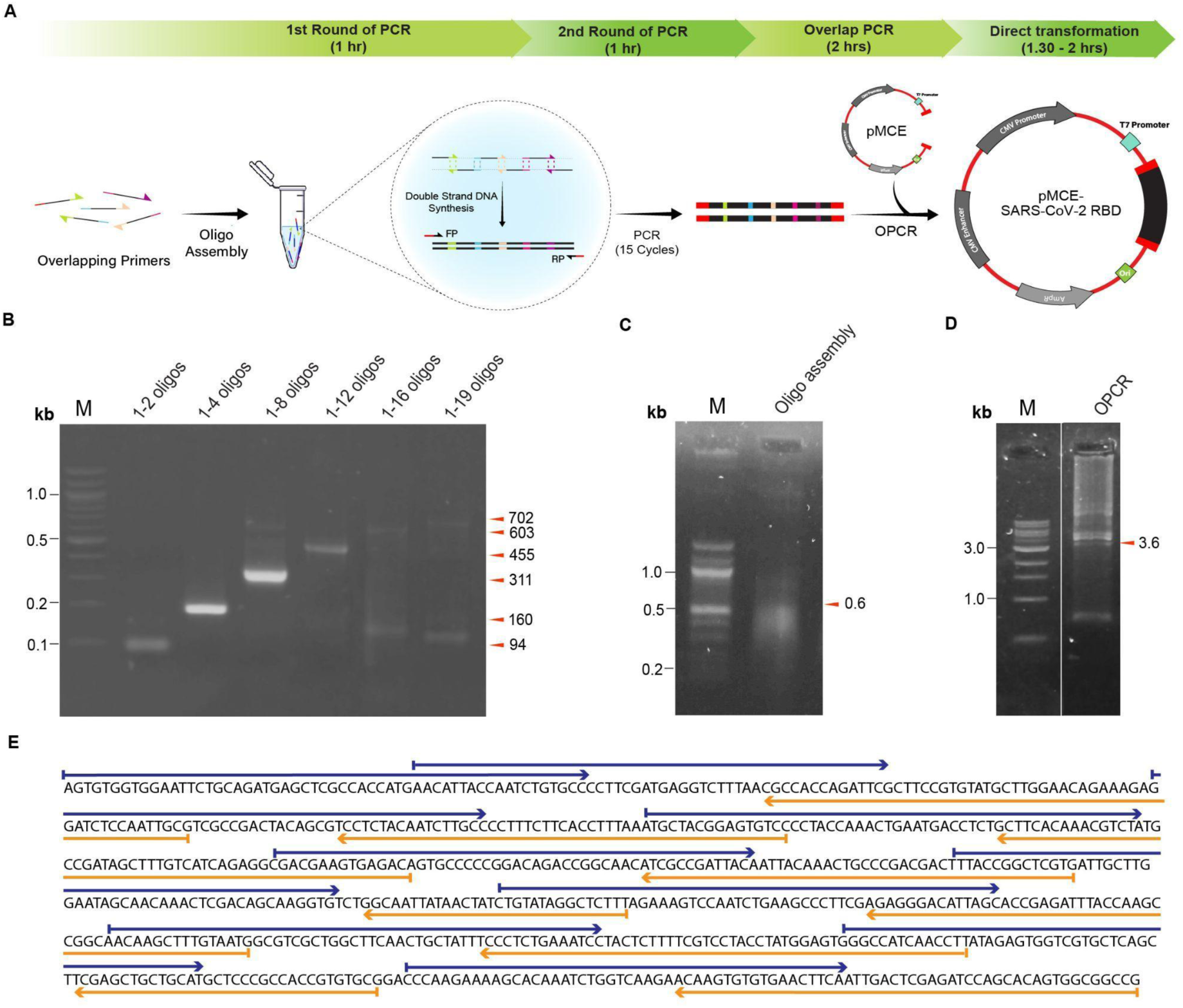
Synthesis of a short gene fragment using O_PCR._ (**A**) Schematic representation of the synthesis of SARS-CoV-2 RBD using oligo-assembly. Agarose gel electrophoresis of (**B**) PCR product of oligonucleotides assembled of sizes ranging from 100 bp to 700 bp; Lane M–100 bp DNA molecular weight marker (**C**) Assembly of oligonucleotides encoding the RBD gene of SARS-CoV-2-Omicron variant; Lane M–100 bp DNA molecular weight marker (**D**) O_PCR_ product of assembled RBD cloned into the pMCE vector; Lane M–1 kb DNA molecular weight marker (**E**) Sanger sequencing result of the assembled RBD gene. Blue and orange arrows indicate forward and reverse primers, respectively. Distinct lanes cropped from one gel image have been assembled as a composite image and separated by white lines and correspond to the marker run on the same gel.

## DISCUSSION

Gene cloning and SDM are fundamental techniques used across the field of life sciences for gene manipulation and functional studies (61–63). However, traditional cloning and mutagenesis methods are often laborious, time-consuming, and require technical expertise (4, 64). Thus, developing rapid, user-friendly strategies to simplify experimental workflows and improve efficiency remains a major focus in molecular biology. While numerous improved methods have recently been introduced to streamline these processes (65–67), high throughput cloning of multiple genes of varying sizes or introducing multiple mutations at distinct loci remains a significant challenge. Here, we present a versatile, PCR-based method adaptable for three key applications: gene cloning, SDM, and short gene synthesis. This approach is based on the principle of O_PCR_ (68) and relies on short homologous regions, typically 15–20 nucleotide sequences between the insert and the plasmid vector (69). These overlapping regions enable efficient annealing and extension by high-fidelity DNA polymerase during the O_PCR_ step, eliminating the need for restriction digestion and ligation (70, 71).

Using this strategy, we initially cloned small gene fragments ranging from 80 bp to 3000 bp into plasmid vectors using a 15 bp overlap and consistently obtained positive clones with high efficiency. While overlaps as short as 12 bp also worked, a 15 bp overlap was found to be optimal, previous studies have shown that 15 to 30 bp is optimal for ligation-independent cloning (72–75). The optimal efficiency likely results from the combined effect of the overlap length and the performance of the high-fidelity polymerase (76, 77). Importantly, shorter overlaps reduce the likelihood of undesirable secondary structures such as hairpins and self-annealing, which can hinder PCR efficiency (78, 79). Cloning of both very short (<200 bp) and large (>7 kb) DNA fragments into plasmids is often challenging (80, 81). Short fragments are often difficult to amplify, purify, and clone due to low yields and degradation during enzymatic digestion or purification steps (82). Moreover, another common problem in cloning is that certain templates (AT or GC-rich) yield very low copy numbers even after multiple attempts of PCR amplification, further purification steps could potentially reduce the DNA concentration, which could hinder downstream applications (83–86). The VSR method overcomes these limitations by eliminating digestion and purification steps entirely, by directly using PCR products for cloning. Moreover, unlike traditional methods that require high DNA concentrations (36, 87), VSR works efficiently with as little as 0.025 to 0.05 pmol of insert and vector, making it ideal for any samples, including low-yield or precious samples (88). Using this method, we successfully cloned gene fragments of various sizes, including very short DNA fragments, with consistently high efficiency. Previous studies have noted that small DNA fragments are difficult to clone due to degradation risks and poor digestion efficiency (82).

We also addressed the challenge of generating large DNA clones (>7 kb) (89), here we showed the efficiency of our method by cloning a 10.3 kb vesicular stomatitis virus (VSV ΔG/GFP) genome and the 32.7 kb adenovirus genome in a short time. These were cloned by amplifying them into smaller fragments ranging from ∼2 kb to ∼5 kb and annealed each fragments along with desired plasmid and complimentary strands were filled by high-fidelity polymerase. Similar approach has also been applied to generate clones of other viruses (90–94). Despite typical PCR-related challenges, such as smearing, degradation, and error-prone amplification (95, 96), we achieved successful cloning of both viral genomes. For the adenovirus genome, high GC content presented amplification difficulties; however, as reported previously (97, 98), these were overcome by adding DMSO and lowering the annealing temperature, leading to improved yield and successful assembly. Furthermore, we demonstrated the ability of this approach to facilitate replacement of a gene in a larger viral genome. The adenoviral fiber gene was replaced with a kanamycin resistance gene *in vitro*, confirming that the method can be readily adapted for precise genome editing applications. Previously, mutations including, insertion, deletion, or substitutions were introduced into adenovirus or other viral genomes by in vivo homologous recombination approach (99–102), our approach eliminates the need for this process.

Similar to gene cloning, SDM is a widely used molecular biology technique for studying gene and protein function (103–106). In this study, we present a simple and robust approach for introducing various types of mutations, including single and multiple nucleotide substitutions, short insertions and deletions, as well as large deletions at multiple loci within a plasmid. Conventionally, SDM requires precise primer design, typically involving 15–20 bp of complementary sequence at the 3′-end of the primer to ensure specific annealing, with the mutation placed toward the 5′-end (107). In traditional protocols, the entire plasmid is amplified using a high-fidelity polymerase with mutagenic primers (forward and reverse), followed by transformation into *E. coli*, where the amplified DNA is circularized (45). These methods also require a pre-existing plasmid clone of the gene of interest and often include additional steps such as DpnI digestion to remove the wild-type template (108). In contrast, our VSR method eliminates many of these constraints. It does not require prior cloning of the gene into a plasmid or the use of specific restriction sites. Instead, mutations are incorporated directly into the primers, which are then used to amplify overlapping DNA fragments from any suitable DNA template, such as plasmid, genomic DNA, PCR product, or cDNA. These fragments are stitched along with the vector using O_PCR_, enabling flexible introduction of various mutations at any desired location within the plasmid. This makes the VSR approach highly adaptable for diverse applications.

A key advantage of our method is its ability to eliminate wild-type DNA contamination, a common issue in standard SDM workflows (87, 109). Since only PCR products with overlapping regions are circularized and successfully transformed, wild-type template DNA is effectively excluded from the final clones, even without DpnI treatment (110). This increases both the specificity and the reliability of the mutagenesis. For example, using this method, we successfully introduced 10 nt substitutions into the SARS-CoV-2 spike gene using primers with only ∼10 bp of 3′ complementarity. Despite the mutations being spread throughout the length of the primers (not confined to the 5′ end), we achieved a mutagenesis efficiency greater than 75%. Furthermore, we applied this method to synthesize a 600 bp gene fragment *de novo* from overlapping oligonucleotides.

In the context of gene cloning and SDM, a common challenge is that a protocol optimized for one gene may not be directly applicable to another due to factors such as gene length, GC content, secondary structures, repeat regions, annealing temperature, or plasmid differences (111, 112). In addition, next-generation sequencing (NGS) technologies have enabled the generation of large sequencing datasets from clinical materials (113, 114), which allows the identification of low-frequency variants during the onset and progression of diseases (115, 116). However, deciphering the functional roles of all these variants remains challenging. Here, we have experimentally validated the adaptability of the O_PCR_-based VSR method to clone 35 different genes ranging from 80 bp to 32.7 kb. These genes were inserted at various sites in different plasmid vectors using either PCR products, cDNA, or restriction-digested DNA as templates. Additionally, we showed that even with relaxed primer design parameters, the method could introduce up to 17 nt substitutions in a single round of mutagenesis. To further evaluate the practicality of the VSR protocol, we tested its compatibility with crude templates, including unpurified PCR products and cDNA. Once the template DNA and plasmid vector are prepared, the entire cloning workflow, including the assembly of single or multiple gene fragments, can be completed within 1.5 to 3 hours, depending on fragment size, followed by transformation. This allows for the generation of recombinant clones in less than 24 hours with high efficiency. Notably, when PCR products derived from plasmid DNA were used directly without purification, we achieved cloning efficiencies of 70–85% using antibiotic selection. Although cloning efficiency from cDNA templates was lower, likely due to contaminants and nonspecific amplification (117), nine genes were successfully cloned, demonstrating the robustness of the method even with suboptimal templates.

Here, we presented a simplified and streamlined approach, capable of cloning a wide range of gene fragments and viral genomes ranging from 80 bp to 33 kb. In addition, the method enabled efficient introduction of diverse site-directed mutations, including substitutions, insertions, and deletions. Moreover, we were able to *de novo* synthesize short gene fragments of size ranging from 90 bp to 700 bp using oligonucleotides. Our method was used to clone 35 distinct genes demonstrating its flexibility. Together, the VSR method is simple and robust approaches that brings cloning, mutagenesis, and de novo gene synthesis in a single platform. This approach could provide an invaluable tool with broader applications in various fields of life sciences.

## MATERIAL AND METHODS

### Preparation of different types of competent cells

#### Calcium chloride method

The preparation of chemical competent cells was performed as described elsewhere (118). Briefly, the primary culture was prepared by inoculating *E. coli* DH5α bacteria from a frozen glycerol stock or picking a single colony from a nutrient agar plate into 5 ml of nutrient broth (NB; HiMedia, #M002). The cells were incubated overnight in a shaking incubator at 37°C with 200 rpm. The following day, 1 ml of the primary culture was inoculated into 100 ml of NB and incubated at 37°C until the OD_600_ reached 0.5-0.6, then the culture was placed on ice for 30 min. All the reagents and plastic wares were kept on ice throughout the procedure. The cells were pelleted at 2500 rpm for 20 min at 4°C, and then gently resuspended in 10 ml of CaCl₂ (100 mM; Sigma, #223506), followed by incubation on ice for 30 min. After a second centrifugation step, the cell pellet was again gently resuspended in 4 ml of CaCl₂. Subsequently, 1.3 ml of chilled 50% glycerol (Sigma, #G6279) was added to the suspension and then divided into 50 µl aliquots in 2 ml microcentrifuge tubes and stored at ‒80°C until further use.

#### Ultracompetent cell (Inoue Method)

The Inoue ultracompetent cells were prepared as described elsewhere (119). Briefly, the primary culture was prepared in a super-optimal broth (SOB) medium as previously described. 1 ml of the primary culture was inoculated into 100 ml of SOB medium and incubated at 18°C with 200 rpm shaking until the OD_600_ reached 0.55, then the culture was placed on ice for 10 min. The cells were harvested by centrifugation and gently resuspended in 8 ml of Inoue buffer (10 mM PIPES (free acid) (Sigma, #P6757), 15 mM CaCl₂·2H₂O, 250 mM potassium chloride (HiMedia, #MB043), 55 mM MnCl₂·4H₂O (HiMedia, #GRM686). The cells were pelleted and resuspended in 2 ml of Inoue transformation buffer. Further, 150 µl of ice-cold DMSO (Sigma, #472301) was added, and incubated on ice for 10 min. 50 µl of the suspension was aliquoted into 2 ml microcentrifuge tubes and snap-frozen by immersing the tubes in a liquid nitrogen bath. The tubes were stored at ‒80°C until further use.

#### Electrocompetent cell

The electrocompetent cells were prepared with minor modifications as described elsewhere (120). The primary culture of *E. coli* DH10B or BJ5183 was prepared as mentioned previously. 5 ml of the primary culture was inoculated into 250 ml of NB at 37 °C, 200 rpm until the OD_600_ reached 0.6-0.8, and then the culture was placed on ice for 30 minutes. The cells were pelleted and resuspended in 35 ml of ice-cold 10% glycerol. This step was repeated twice using 10 ml and 5 ml of 10% glycerol, respectively. The cell pellets were resuspended in 1 ml of 10% glycerol, and the cell density at 1:100 dilution was determined at OD_600_ (OD_600_ = 0.8 corresponds to ∼2.5×10^8 cells/ml per vial). Once the OD was set up to 0.8, it was aliquoted as 50 µl in 2 ml microcentrifuge tubes. It is preferable to prepare fresh electrocompetent cells before transformation instead of storing the cells at –80 °C.

### Generation of a mini-cloning plasmid DNA (pMCE)

To generate the pMCE vector, sequences containing minimal essential regulatory elements, including the origin of replication (ori), cytomegalovirus (CMV) and bacteriophage T7 promoter, bovine growth hormone (bGH) polyadenylation (polyA) signal, and ampicillin resistance gene (amp^R^), were PCR amplified from the pcDNA3.1+ expression vectors using PfuUltra II Fusion High-fidelity Polymerase (Agilent, #600672) following the manufacturer’s instructions. The PCR product was purified using the FavorPrep GEL/PCR Purification Mini Kit (Favorgen Biotech Corp, #FAGCK001-1). 100 ng of the purified PCR product was circularized using 1 µl (400 units) of T4 DNA Ligase (NEB, #M0202S) for 1 h at room temperature. Then, 1 µl of the ligated product was transformed into chemically competent cells of *E. coli* DH5α following the heat shock method and incubated for 12–16 h at 37°C. Further, the colonies were screened for positive clones by colony PCR (as mentioned below) using CMV forward primer 5’-CGCAAATGGGCGGTAGGCGTG-3’ and bGH reverse primer 5’-CACCTTCCAGGGTCAAGGAA-3’, and the plasmid was further confirmed using restriction digestion and Sanger sequencing. Then the plasmid was linearized using the desired restriction enzymes and used for downstream applications of gene cloning, SDM, and gene synthesis.

### VSR Method

The VSR method for cloning, mutagenesis, and oligo assembly works on three simple steps:

1. Primer design and preparation of the DNA template(s) for cloning, site-directed mutagenesis, and gene synthesis.
2. Cloning of the DNA template into a plasmid using O_PCR._
3. Bacterial transformation, screening, and validation of clones using colony PCR and sequencing.

### 1. Primer design and preparation of the DNA template(s) for cloning, site-directed mutagenesis, and gene synthesis

#### Primer Design for Gene cloning

All primers used in this study were purchased from Eurofins Genomics (India). For cloning a gene of interest, a pair of forward and reverse primers were designed as follows. Each primer contained two components: (i) a 20 nt gene-specific sequence (GSS) at the 3′ end and (ii) a 15 nt sequence complementary to the vector (OS), which generates an overhang during amplification of the gene of interest (Fig. 1). The resulting PCR products enable directional cloning of the desired gene. All cloning primers in this study were designed using this format (C-FP: 5′-15OS+20GSS-3′ and C-RP: 5′-15OS+20GSS-3′) and are hereafter referred to as *overlapping cloning primers*. All the oligonucleotides used for cloning in this study is included in Supplementary Table S1-S3. Additionally, in certain cases, a restriction site(s) or a short tag sequence was incorporated immediately adjacent to the gene-specific region (5′-15OS+NNNN+20GSS-3′). For cloning multiple fragments, 15 bases complementary to the sequence of the adjoining fragment were added at the 5′ end of each primer.

#### Primer Design for SDM

To introduce mutations at specific sites, we employed multiple fragment cloning strategy as described above. Instead of modifying plasmid DNA directly, the desired mutations were incorporated into oligonucleotide primers, which were then used to amplify overlapping fragments from gDNA or cDNA. Two sets of primers were designed (Supplementary Fig. S1): the first set consisted of a cloning forward primer (C-FP) and a mutagenic reverse primer (M-RP), and the second set consisted of a mutagenic forward primer (M-FP) and a cloning reverse primer (C-RP). The forward primer of the first set and the reverse primer of the second set contained a gene-specific sequence followed by a 15 nt overlap complementary to the cloning vector. While the reverse primer of the first set and the forward primer of the second set carrying the mutations were complementary to each other. Mutagenic primers consisted of 10–20 nt complementary to the target sequence at the 3′ end, with the desired mutation (substitution, short insertion, or deletion) and a 15 nt overlap incorporated at the 5′ end. Both mutagenic primers shared a 15 nt overlap at their 5′ ends to facilitate fragment joining. Note that all our mutagenesis primers were designed in the same manner (5′-15OS+Ms+20GSS-3’). These will henceforth be referred to as *overlapping mutagenesis primers*. For introducing or deleting short nucleotide sequences in a gene, the primers were designed similar to the overlapping mutagenesis primers, consisting of a gene-specific sequence followed by the desired insertion/deletion and subsequently overlapping sequence (5’-15OS+IN+20GSS-3’/ 5’-15OS+DEL+20GSS-3’). For large deletion mutagenesis, mutagenic primers were designed such that the 3′ end contained 20 nt complementary to the sequence immediately upstream of the deletion site, while the 5′ end contained 15 nt complementary to the sequence immediately downstream (Supplementary Fig. S2). The summary of the oligonucleotides used for introducing diverse mutations is given in Supplementary Table S4.

#### Primer Design for Gene synthesis

To synthesize the receptor binding domain (RBD) of the SARS-CoV-2 Omicron variant (GenBank: MN908947), 600 bp in size, the nucleotide sequence of codon-optimized SARS-CoV-2 WT was used as a reference, and the primers were designed based on the mutations reported in the literature. Eighteen overlapping oligonucleotides, each of ∼50 bp length, were designed with 15–22 bp of unique overlap between the adjacent ones adjusted to a predicted Tm of 50°C or greater. The primers were arranged in alternating forward (sense) and reverse (antisense) orientations such that each adjacent pair overlaps (Supplementary Fig. S3). The first and last primers encoding the start and stop of the gene fragment, respectively, have 15 bp of overlap with pMCE to facilitate cloning. The secondary structures of the primers were predicted using the OligoCalc online tool. Further, the primers were corrected to have as minimal secondary structure formation as possible. The summary of the oligonucleotide set with detailed information is provided in the Supplementary Table S5.

### Preparation of DNA template

Template preparation is a critical step in any cloning process. In our method, the desired DNA fragments ranging from 80 bp to 5 kb were PCR-amplified from either genomic DNA or a cDNA library using overlapping cloning primers (Supplementary Table S1). The PCR products were then used for cloning into a vector, with or without any further purification.

For larger genes (>7 kb) or those difficult to amplify in a single reaction, amplification was performed as shorter overlapping fragments (0.1−5.8 kb each), which were subsequently assembled into a single continuous DNA fragment suitable for cloning. For successful assembly, each fragment was designed with 15 bp overlaps for smaller fragments and 40−100 bp overlaps for larger fragments at the ends. (Supplementary Tables S2, S3). These overlaps facilitate specific hybridization between neighbouring fragments during annealing, which are then extended by the DNA polymerase and amplified into a continuous DNA sequence. The terminal ends of the assembled gene also contain sequences complementary to the cloning vector, allowing direct cloning into a plasmid vector.

#### Preparation of cDNA library

RNA samples from various mammalian cell lines, including human embryonic kidney 293T cells (HEK293T-ATCC CRL-3216, Manassas, VA, USA), the human liver cell line, and the human lung cell line (Huh7 and A549 (Erasmus Medical Centre, Rotterdam, Netherlands)), were isolated using the RNeasy Mini Kit (Qiagen, #74104) as per the manufacturer’s instructions. To synthesize the cDNA library, 0.5‒1 µg of the isolated RNA was added to the reaction mix containing 0.5 µl (40 U/µl) of RNase inhibitor (Promega, #N2618), 1 µl (2.5 mM each) of dNTP (Takara, #4026), and 0.5 µl of random hexamer primers (10 µM). The reaction mix was heated at 65°C for 10 min and quickly chilled at 4°C. Further, the reaction mixture was collected by brief centrifugation and added to the reaction mix containing 1 µl of 5× M-MLV RT buffer, 1 µl (0.1 M) of DTT (Promega, #00677079), and 1 µl (200 U/µl) of M-MLV reverse transcriptase (Promega, #M1708) made up to 25 µl using nuclease-free water (Promega, #P1199). The reaction mix was incubated at 25°C for 10 min followed by 37°C for 1 hour. Finally, the reaction was heated at 70°C for 10 minutes to inactivate the enzyme. The cDNA template was used for PCR amplification of specific genes using overlapping primers.

#### PCR amplification

The PCR master mix was prepared on ice with the corresponding forward and reverse primers. The mix was prepared using the following components: 10 µl (10×) of Primestar GXL Polymerase buffer, 4 µl (200 mM) of dNTP mix, 1 µl (10 µM) of forward and reverse primers, 1 µl (1.25 U/µl) of Primestar GXL DNA Polymerase (Takara, #R050A), and 10-50 ng of DNA template. The total volume was adjusted to 50 µl with nuclease-free water. Further, the PCR mixture was incubated at 98°C for 30 s, followed by 30 cycles of denaturation at 98°C for 20 s, annealing at 55°C for 20 s, and extension at 68°C for 1 min per kb, followed by a final extension at 68°C for 5 min. 5 µl of the PCR product was run on a standard agarose gel electrophoresis. Upon observation of a single band of the desired size during analysis, the PCR product was used with or without PCR purification. If several bands were detected, the predicted agarose gel size was excised and subjected to gel purification. The DNA molecular weight markers (100 bp and 1 kb) are purchased from NEB.

#### Preparation of vector

The desired plasmids were linearized with restriction enzymes as per the manufacturer’s protocol (New England Biolabs), generating either blunt or cohesive ends. To eliminate the undigested plasmids, the digested product was analyzed on standard agarose gel electrophoresis, and the linearized plasmid band was excised from the gel, purified using the gel/PCR purification kit, and subsequently used for performing cloning or mutagenesis.

### Cloning of DNA Fragments into a Vector Using OPCR

To clone the PCR amplified fragment into plasmid DNA by O_PCR,_ the PCR master mix was prepared by adding 0.025‒0.05 pmol of desired digested vector and purified insert DNA or 25 ng of digested vector and 1‒2 µl of direct PCR product template to master mix consisting of 5 µl (10×) of Primestar GXL Polymerase buffer, 2 µl (200 mM) of dNTP mix, 0.5 µl (1.25 U/µl) of Primestar GXL DNA polymerase adjusted to 25 µl using nuclease free water. The reaction condition of initial denaturation at 98°C for 30 s, followed by 12-15 cycles of denaturation at 98°C for 10 s, annealing at 55°C for 20 s, and extension at 68°C for 10 min, followed by a final extension at 68°C for 10 min was used to allow the annealing of gene fragment to the digested vector.

### Bacterial transformation, screening, and selection of clones using colony PCR

#### Heat shock transformation

One microliter of the O_PCR_ product was directly added to the *E. coli* DH5α competent cells and gently tapped to ensure even distribution of the DNA. The mix was further incubated on ice for 30 min. The cells were subjected to heat shock at 42°C for 45 s in a water bath, followed by incubation on ice for 2 min. Further, 450 µl of pre-warmed NB medium was added to the tubes and placed in a shaking incubator at 200 rpm at 37°C for 1 h (approximately three cell division cycles). 100 µl of the transformation reaction was spread on a warm NB agar plate containing the appropriate antibiotics and incubated at 37°C overnight. After incubation, at least 10 colonies were picked for screening positive clones using colony PCR.

#### Electroporation-mediated transformation

Prior to electroporation, the cuvettes (Biorad, #1652082) were pre-chilled, and the recovery NB media without antibiotics and NB plates with selective antibiotics were pre-warmed at 37°C. The electrocompetent cells were thawed on ice for 10 minutes, and 1 µl of the O_PCR_ product was added to the cells. The mix was gently transferred without introducing any bubbles into a pre-chilled cuvette, ensuring that the cells settled at the bottom of the cuvette (0.1 cm). The cells were electroporated using the EC-1 program (1.8 kV (E= 18 kV/cm) in a Micropulser electroporator (Biorad). Immediately after the electroporation, 1 ml of pre-warmed NB media was added to the cuvette, gently mixed, and transferred to a fresh 2 ml microcentrifuge tube. The cells were incubated at 37°C and 200 rpm for 1 hour, and subsequently, 100-200 µl of the culture was plated onto the NB agar plate supplemented with a selective antibiotic. The plates were incubated at 37°C for 16-28 hours, and colonies were screened using colony PCR.

#### Colony PCR

The individual bacterial colonies were picked using a toothpick and mixed in 10 µl of NB media without any antibiotics. 1 µl of the bacterial culture was added to the PCR mix, which consists of 1 µl (10×) of Taq DNA Polymerase buffer, 0.4 µl (200 mM) of dNTP mix, 1U of Taq DNA Polymerase (NEB, M0273S), and 0.3 µl (10 µM) of forward and reverse primers each adjusted to a volume of 10 µl using nuclease-free water. The PCR mixture was mixed gently, and PCR was performed in a thermocycler using the following conditions: initial denaturation at 92°C for 10 min. Repeat the following process for 25 cycles: denaturation at 92°C for 20 s, annealing at 55°C for 20 s, extension at 72°C for 1 min/kb, and final extension at 72°C for 5 min. The products were analyzed using agarose gel electrophoresis, and images were captured using a ChemiDoc system.

#### Sanger sequencing

The templates, either a PCR product or plasmid DNA, were purified using column purification to achieve a purity of 1.8. Either 100 ng of PCR product or 200 ng of plasmid DNA template was added to the reaction mix consisting of 0.5 µl of BigDye Terminator 3.1 ready reaction mix (Applied Biosystems, #4336697), 1.75 µl of BigDye Terminator v1.1 & v3.1 5x sequencing buffer, and 1 µl (10 µM) of forward primer or reverse primer, adjusted to 10 µl using nuclease-free water. The reaction mix was incubated for initial denaturation at 96°C for 1 min, followed by 25 cycles of 96°C for 10 s, 50°C for 0.5 s, and 60°C for 4 min, with each condition adjusted to a ramp rate of 1°C/s. Further, to remove the remaining dye terminators, primers, and buffer, the product was purified using ethanol/EDTA precipitation or plate and spin column purification and used for capillary electrophoresis.

### Statistical Analysis

All graphs presented in the paper were analyzed using GraphPad Prism 9.4.1. A two-tailed, unpaired t-test was performed to calculate the statistical significance between blunt and cohesive end cloning efficiency. Bar graphs are expressed as the mean, and each point represents individual genes. Significance value *** indicates P-value < 0.01. The before-and-after graphs were not evaluated for any significance test since their aim was to identify the correlation between the insert size and cloning efficiency within the same sample.

## Supporting information

Kavitha_VSR Method_Supplementary Tables_Figures_Biorxiv

## ACKNOWLEDGEMENTS

The methods were tested by multiple research scholars in our laboratory. We gratefully thank Karthika T, Malavika Prakash, Harshal Srivastava, Saran M, Shreosi Basu, and Azeem Abdul Kalam from Virology Scientific Research lab and Tejas T. M. from HapGen lab, School of Biology, IISER-TVM, for their contribution in generating gene clones and site directed mutagenesis using our method.

## AUTHOR CONTRIBUTIONS

Conceptualization: V.S.R. Methodology: V.S.R., K.M.S., G.K., A.D.V.R. Investigation: V.S.R., K.M.S., G.K., J.S., A.D.V.R., N.B., A.P.S. Visualization: K.M.S., G.K., J.S., A.D.V.R., N.B Supervision: V.S.R. Writing: V.S.R, K.M.S. Writing-review & editing: V.S.R., K.M.S., G.K., A.K.D.R.

## DATA AVAILABILITY

All the data supporting this article are included in the main text and its online supplementary material.

## SUPPLEMENTARY DATA

Supplementary Data are available online.

