## Supplementary material for "A Versatile, Simple, and Rapid (VSR) method for generating gene clones, diverse mutants, and short gene fragments for molecular biology applications": Kavitha_VSR Method_Supplementary Tables_Figures_Biorxiv

Kavitha Modenkattil Sethumadhavan[a], Gayatri Kandpal[a], Jyothilakshmi Sajimon[a], V. R. Akshay Das[a], Nirmegh Basu[a], Athulya Prakash Simla[a], V. Stalin Raj[a]*
[a]Virology Scientific Research (VSR) Laboratory, School of Biology, Indian Institute of Science Education and Research, Thiruvananthapuram, Kerala, 695551, India

### Supplementary Material

### 1. Supplementary tables

#### Table S1. List of overlapping primers used in this study to perform gene cloning

| S.No | Name of Gene/Fragment | Primer Code | Primer Sequence (5' -> 3') | Number of overlapping bases | GC % of overlapping region | Ends |
|---|---|---|---|---|---|---|
| 1 | IL4 | CEC-1-FP | GGGTCGCGGATCCGAATTCCATAACTTCAGTATTGCCAT | 13 | 35 | Cohesive |
| | | CEC-1-RP | GGTGGTGGTGCTCGAGTGCGGCCGCATGCCTGTAGTATTTCTGTTG | 17 | 38 | |
| 2 | Ferret γ chain | CEC-2-FP | GTAGGCCTCGTACGCTTAATTAAACCATGGTCCCCGCAGTGGTCTTG | 15 | 56 | Cohesive |
| | | CEC-2-RP | GCGGAATTCCGGATCCTCACTTATCGTCGTCATCCTTGTAATCCTCGAGCCTAGGTTACTGGGATG | 16 | 51 | |
| 3 | IL2 | CEC-3-FP | GGGTCGCGGATCCGAATTCGCACCTACTACTTCAAGCTC | 13 | 50 | Cohesive |
| | | CEC-3-RP | GGTGGTGGTGCTCGAGTGCGGCCGCGATGCTTTGACAAAAAGTAATC | 17 | 32 | |
| 4 | IFN-A | CEC-4-FP | GGGTCGCGGATCCGAATTCTGTGACCTGCCTCAGGACCAC | 13 | 62 | Cohesive |
| | | CEC-4-RP | GGTGGTGGTGCTCGAGTGCGGCCGCCTTCCTGCTCCGCAATCTC | 17 | 58 | |

| | | | | | | |
|---|---|---|---|---|---|---|
| 5 | IFN-B | CEC-5-FP | GGGTCGCGGATCCGAATTCATGAACTATAACTTACTTCG | 13 | 30 | Cohesive |
| | | CEC-5-RP | GGTGGTGGTGCTCGAGTGCGGCCGCGTTAGGGAGAGAATCTGTAAG | 17 | 43 | |
| 6 | ARF6 | CEC-6-FP | GCTCCGGACTCAGATCTCGAGGCGGGAAGGTGCTATCCAAA | 15 | 50 | Cohesive |
| | | CEC-6-RP | GGATCCCGGGCCCGCGGTACCTTAAGATTTGTAGTTAGAGGT | 15 | 33 | |
| 7 | Caveolin | CEC-7-FP | GCTCCGGACTCAGATCTCGAGGCTCTGGGGGCAAATACGTA | 15 | 50 | Cohesive |
| | | CEC-7-RP | GGATCCCGGGCCCGCGGTACCTTATATTTCTTTCTGCAAGTT | 15 | 24 | |
| 8 | eGFP | CEC-8-FP | GGAATTGATCCGCGGCCGCAGCCACCATGGTGAGCAAGGG | 20 | 65 | Cohesive |
| | | CEC-8-RP | GGAATTCCGGATCCGTTAATTTACTTGTACAGCTCGTCCA | 19 | 55 | |
| 9 | Clathrin | CEC-9-FP | GCTCCGGACTCAGATCTCGAGGCGCTGATGACTTTGGCTTC | 15 | 50 | Cohesive |
| | | CEC-9-RP | GGATCCCGGGCCCGCGGTACCCTAGCGGGACAGTGGCGTCTG | 15 | 72 | |
| 10 | Human FcGRIII | CEC-10-FP | CCTGCAGGAATTGATCCGCGGCCGCAACATGTGGCAGCTGCTCCTC | 17 | 58 | Cohesive |
| | | CEC-10-RP | GTGATGGTGATGATGCTCGAGTTTGTCTTGAGGGTCCTTTC | 15 | 62 | |
| 11 | sCAR, eGFP 1-10 | CEC-11-FP | CAGGAATTGATCCGCGGCCGCGCCACCATGTTGAGTATCACTACTCCTGA | 21 | 48 | Cohesive |
| | | CEC-11-RP | CTCACCATGGTGGCACCGGTTGATCCACCTCCACCAGATC | 20 | 55 | |
| 12 | SARS CoV-2 S1 | BEC-12-FP | TGTGGTGGAATTCTGCAGATGAGCTCCAATGCGTCAACCTCACAAC | 20 | 50 | Blunt |
| | | BEC-12-RP | CGGCCGCCACTGTGCTGGATCTCGAGGTCTTCTAGGGGAGTTGGTCTG | 20 | 55 | |
| 13 | NRP1 | BEC-13-FP | TGTGGTGGAATTCTGCAGATACCGGTGCCACCATGGAGAGGGGGCTGCCGCT | 20 | 72 | Blunt |
| | | BEC-13-RP | CGGCCGCCACTGTGCTGGATCCTAGGTTATGCCTCCGAATAAGTACTCTGTG | 20 | 20 | |
| 14 | mGluR2 | BEC-14-FP | TGGAATTCTGCAGATACCGGTGCCACCATGGGATCGCTGCTTGCGCT | 15 | 70 | Blunt |

| | | | | | | |
|---|---|---|---|---|---|---|
| | | BEC-14-RP | GCCACTGTGCTGGATCCTAGGTTAAAGCGATGACGTTGTCGAGT | 15 | 67 | |
| 15 | DelCyt NRP1 | BEC-15-FP | TGTGGTGGAATTCTGCAGATACCGGTGCCACCATGGAGAGGGGGCTGCCGCT | 20 | 70 | Blunt |
| | | BEC-15-RP | CGGCCGCCACTGTGCTGGATCCTAGGTTACAGCACGACCCCACAGACAG | 20 | 67 | |
| 16 | SARS CoV-2 S | BEC-16-FP | TGTGGTGGAATTCTGCAGATGAGCTCCAATGCGTCAACCTCACAAC | 20 | 50 | Blunt |
| | | BEC-16-RP | CGGCCGCCACTGTGCTGGATCTCGAGTCAGCAGCAGCTTCCGCAGC | 20 | 65 | |
| 17 | L gene | | | | | |
| | Fragment-1-LA | BEC-17-FP | GGACCCTGTGTTCTCACTGT | - | - | Blunt |
| | | BEC-17-RP | AAAGCGCGCGCTCTCTTCATTCCATTTAGATTGTGGAGAA | 40 | -- | |
| | Fragment-2-RA | BEC-18-FP | TTCTCCACAATCTAAATGGAGGATGAATGTCAGCTACTGG | 40 | - | |
| | | BEC-18-RP | ACATAACACAAACGATTCTTCAGAAGAACTCGTCAAGAA | 39 | - | |
| | Fragment-3- Kan | BEC-19-FP | ATGTTGGAGAAAATCCATGAAGAATCGTTTGTGTTATGTT | 40 | - | |
| | | BEC-19-RP | CCTAAGAATATGTCTGTTACCC | - | - | |
| 18 | NP Fragment | | | | | |
| | Fragment-1NP | BEC-20-FP | AGTGTGGTGGAATTCTGCAGATTAATACGACTCACTATAGGGA | 22 | | Blunt |
| | | BEC-21-RP | TGGAGTCTCCTCATGATTTTCTACAGAGAATATTTGACTC | - | - | |
| | Fragment-2 Trailer | BEC-22-FP | GAGTCAAATATTCTCTGTAGAAAATCATGAGGAGACTCCA | - | - | |
| | | BEC-22-RP | TCAAAATCGTGGACTTCCATAAAGTTTCTCCTGAGCCTTT | - | - | |
| | Fragment-3 ribozyme | BEC-23-FP | TATCTGGTTTTGTGGTCTTCGTGGCCGGCATGGTCCCAGCCT | - | - | |
| | | BEC-23-RP | AGGCTGGGACCATGCCGGCCACGAAGACCACAAAACCAGA | - | - | |

| | | | | | | |
|---|---|---|---|---|---|---|
| 19 | Fragment-1-eGFP11 | CEC-24-FP | GGAGACCCAAGCTGGGCCACCATGCGCGATCACAT | 15 | | Cohesive |
| | | CEC-24-RP | GTCACAAGATTTGGGGGATCCCGGGCCCGCGGTACC | 15 | | |
| | Fragment-2 IgG-Fc Tag | CEC-25-FP | CCCAAATCTTGTGACAAAAC | 15 | | |
| | | CEC-25-RP | ACGGGCCCTCTAGACTCATTTACCCGGAGACAGGG | 14 | | |
| 20 | Fragment-1 Leader | BEC-26-FP | TAATACGACTCACTATAGGGACGAAGACAAACAAACCATT | 22 | | Blunt |
| | | BEC-26-RP | TCAAAATCGTGGACTTCCATAAAGTTTCTCCTGAGCCTTT | 40 | | |
| | Fragment-2 L gene1 | BEC-27-FP | AAAGGCTCAGGAGAAACTTTATGGAAGTCCACGATTTTGA | - | - | |
| | | BEC-27-RP | TCGCTGGACTCATTCCCATA | - | - | |
| | Fragment-3 L gene2 | BEC-28-FP | TCCAACCTCTCTGAACATCG | - | - | |
| | | BEC-28-RP | TTGGAGGCATCTCTAGCATC | - | - | |
| | Fragment-4 L gene3 | BEC-29-FP | TACGACCACCCCTTACCCAA | - | - | |
| | | BEC-29-RP | AGGCTGGGACCATGCCGGCCACGAAGACCACAAAACCAGA | - | - | |
| | Fragment-5 ribozyme | BEC-30-FP | TATCTGGTTTTGTGGTCTTCGTGGCCGGCATGGTCCCAGCCT | 42 | | |
| | | BEC-30-RP | AGGCTGGGACCATGCCGGCCACGAAGACCACAAAACCAGA | 40 | | |
| 21 | IL-4 | CEC-1-FP | GGGTCGCGGATCCGAATTCCATAACTTCAGTATTGCCAT | 13 | 35 | Cohesive |
| | | CEC-1-RP | GGTGGTGGTGCTCGAGTGCGGCCGCATGCCTGTAGTATTTCTGTTG | 17 | 38 | |
| 22 | IFN-alpha | CEC-4-FP | GGGTCGCGGATCCGAATTCTGTGACCTGCCTCAGGACCAC | 13 | 62 | Cohesive |

| | | | | | | |
|---|---|---|---|---|---|---|
| | | CEC-4-RP | GGTGGTGGTGCTCGAGTGCGGCCGCCTTCCTGCTCCGCAATCTC | 17 | 58 | |
| 23 | IFN beta | CEC-5-FP | GGGTCGCGGATCCGAATTCATGAACTATAACTTACTTCG | 13 | 63 | Cohesive |
| | | CEC-5-RP | GGTGGTGGTGCTCGAGTGCGGCCGCGTTAGGGAGAGAATCTGTAAG | 17 | 68 | |
| 24 | eGFP | BEC-31-FP | TGTGGTGGAATTCTGCAGATACCGGTCGCCACCATGGTGA | 20 | 65 | Blunt |
| | | BEC-31-RP | AGCGGCCGCCACTGTGCTGGATGCGGCCGCTTTACTTGTACA | 22 | 55 | |
| 25 | Tetraspanin (CD9) | BEC-32-FP | TGGAATTCTGCAGATGGTACCACCATGCCGGTCAAAGGAGGC | 15 | 44 | Blunt |
| | | BEC-32-RP | GCCACTGTGCTGGATCTCGAGCTAGACCATCTCGCGGTT | 15 | 53 | |
| 26 | CD63 | BEC-33-FP | TGGAATTCTGCAGATACCGGTACCATGGCGGTGGAAGGAGGAAT | 15 | 55 | Blunt |
| | | BEC-33-RP | GCCACTGTGCTGGATCCTAGGCTACATCACCTCGTAGCCAC | 15 | 55 | |
| 27 | CD80 | BEC-34-FP | TGGAATTCTGCAGATGAATTCACCATGGGCCACACACGGAGGCA | 15 | 65 | Blunt |
| | | BEC-34-RP | GCCACTGTGCTGGATGCTAGCTTATACAGGGCGTACACTTT | 15 | 47 | |
| 28 | HVEM | BEC-35-FP | TGGAATTCTGCAGATACCGGTACCATGGAGCCTCCTGGAGACTG | 15 | 60 | Blunt |
| | | BEC-35-RP | GCCACTGTGCTGGATTTAATTAATCAGTGGTTTGGGCTCCTCCCCG | 15 | 70 | |
| 29 | CD4 | BEC-36-FP | TGGAATTCTGCAGATGAGCTCACCATGGATTATCAAGTGTCAAG | 15 | 35 | Blunt |
| | | BEC-36-RP | GCCACTGTGCTGGATCTCGAGTCACAAGCCCACAGATATTT | 15 | 40 | |
| 30 | Ephrin B2 | BEC-37-FP | TGGAATTCTGCAGATGGTACCACCATGGCTGTGAGAAGGGACTC | 15 | 55 | Blunt |
| | | BEC-37-RP | GCCACTGTGCTGGATCCCGGGTCAGACCTTGTAGTAAATGT | 15 | 35 | |
| 31 | Cathepsin L | BEC-38-FP | TGGAATTCTGCAGATGCTAGCACCATGAATCCTACACTCATC | 15 | 40 | Blunt |
| | | BEC-38-RP | GCCACTGTGCTGGATCTCGAGTCACACAGTGGGGTAGCT | 15 | 53 | |

| | | | | | | |
|---|---|---|---|---|---|---|
| 32 | AGTR2 | BEC-39-FP | TGGAATTCTGCAGATACCGGTACCATGGCGTGGCGGTGCCCCAG | 15 | 55 | Blunt |
| | | BEC-39-RP | GCCACTGTGCTGGATCCTAGGTTAAGACACAAAGGTCTCCATTT | 15 | 40 | |
| 33 | KIM1 | BEC-40-FP | TGGAATTCTGCAGATGGATCCACCATGCATCCTCAAGTGGTC | 15 | 40 | Blunt |
| | | BEC-40-RP | GCCACTGTGCTGGATCTCGAGTTAGTCCGTGGCATAAAG | 15 | 53 | |
| 34 | CD147 | BEC-41-FP | TGGAATTCTGCAGATTTAATTAAACCATGGCGGCTGCGCTGTTCGT | 15 | 65 | Blunt |
| | | BEC-41-RP | GCCACTGTGCTGGATCCTAGGTTAGGAAGAGTTCCTCTGGCGGA | 15 | 60 | |
| 35 | CD71 | BEC-42-FP | TGGAATTCTGCAGATACCGGTACCATGATGGATCAAGCTAGATC | 15 | 40 | Blunt |
| | | BEC-42-RP | GCCACTGTGCTGGATTTAATTAATCAAAACTCATTGTCAATGTCCC | 15 | 40 | |
| 36 | DAG-1 | BEC-43-FP | TGGAATTCTGCAGATGAATTCACCATGAGGATGTCTGTGGGCCT | 15 | 55 | Blunt |
| | | BEC-43-RP | GCCACTGTGCTGGATGCTAGCTTAAGGTGGGACATAGGGAG | 15 | 60 | |

**Blue**: Overlapping region, **Red**: Template-specific region, **Black**: Restriction sites and/or Kozak sequence, **Green**: 6X Histidine Tag sequence

### Table S2. List of overlapping primers used for cloning VSV ΔG/GFP genome

| Primer Code | Primer sequences (5' - 3') |
|---|---|
| VSV-F1-1-FP | AGTGTGGTGGAATTCTGCAGATTAATACGACTCACTATAGGGA |
| VSV-F1-2-FP | TAATACGACTCACTATAGGGACGAAGACAAACAAACCATT |
| VSV-F1-RP | TCATTTGAAGTGGCTGATAGAATCCAGGA |

| | |
|---|---|
| **VSV-F2-FP** | TCCTGGATTCTATCAGCCACTTCAAATGA |
| **VSV-F2-RP** | TCGGTCTCAAAATCGTGGACTTCCAT |
| **VSV-F3-FP** | ATGGAAGTCCACGATTTTGAGACCGA |
| **VSV-F3-RP** | TCGCTGGACTCATTCCCATA |
| **VSV-F4-FP** | TCCAACCTCTCTGAACATCG |
| **VSV-F4-RP** | TTGGAGGCATCTCTAGCATC |
| **VSV-F5-FP** | TACGACCACCCCTTACCCAA |
| **VSV-F5-RP** | AGGCTGGGACCATGCCGGCCACGAAGACCACAAAACCAGA |
| **VSV-F6-HDVr-FP** | TATCTGGTTTTGTGGTCTTCGTGGCCGGCATGGTCCCAGCCT |
| **VSV-F5-HDVr-RP** | TCGAGCGGCCGCCACTGTGCTGGATTTTCGGGCTTTGTTAGCAGC |

### Table S3. List of overlapping primers used for cloning Ad5 genome

| Primer Code | Primer sequences (5' - 3') |
|---|---|
| **AdV-F1-FP** | GTAGTCATGCTCATGCAGATAAAGG |
| **AdV-F1-RP** | GACAAATTTCCCAAGTGCACCGGAT |
| **AdV-F2-FP** | GTATTTTTAGGTTGGCTATGTTCCC |
| **AdV-F2-RP** | CTCTACCGCCAGCCGCCGCCGCACT |
| **AdV-F3-FP** | TGAGACGCGAGTAAGCCCTCGAGTC |
| **AdV-F3-RP** | GCAGGTGCGGCGTCTGGCGTCAGTA |
| **AdV-F4-FP** | CGTTCCTGCTCTCACAGATCACGG |
| **AdV-F4-RP** | GCAGGTGCGGCGTCTGGCGTCAGTA |
| **AdV-F5-FP** | GGCTTCTATATCCCAGAGAGCTACA |
| **AdV-F5-RP** | CGGTAAGCTCCGCATTTGGCGGTCTAGATTGGTCTTCGTAGAACCTAA |
| **AdV-F6-Kan-FP** | TCTACGAAGACCAATCTAGACCGCCAAATGCGGAGCTTAC |
| **AdV-F6-Kan-RP** | GTTTATGCAGAAACCCGCAGACATG |

### Table S4. List of overlapping primers used for introducing site-directed mutagenesis

| S. No | Mutation Type | Name of the Gene | Primer Code | Nucleotide change | Mutagenic Primer sequences (5' - 3') | Insert size |
|---|---|---|---|---|---|---|
| 1 | Single/Point | SARS CoV-2 S1 | BEC-SDM-1-FP | | **TGTGGTGGAATTCTGCAGATGAGCTCCAATGCGTCAACCTCACAAC** | 2 kb |
| | | | SDM-1-FP | | **AGCTTTGTCATCAGTGGCGACGAAGTGAGACAGAT** | |

| | | | | | | |
|---|---|---|---|---|---|---|
| | | | SDM-2-RP | 1 nt substitution | TCTCACTTCGTCGCCACTGATGACAAAGCTATCGGCA | |
| | | | BEC-SDM-2-RP | | CGGCCGCCACTGTGCTGGATCTCGAGGTCTTCTAGGGGAGTTGGTCTG | |
| 2 | Single/Point | SARS-CoV-2 complete spike | BEC-SDM-1-FP | | TGTGGTGGAATTCTGCAGATGAGCTCCAATGCGTCAACCTCACAAC | 4 kb |
| | | | SDM-1-FP | 1 nt substitution | AGCTTTGTCATCAGTGGCGACGAAGTGAGACAGAT | |
| | | | SDM-2-RP | | TCTCACTTCGTCGCCACTGATGACAAAGCTATCGGCA | |
| | | | BEC-SDM-3-FP | | CGGCCGCCACTGTGCTGGATCTCGAGTCAATTGAAGTTCACACACTTGT | |
| 3 | Single aa insertion | SARS CoV-2 S1 | BEC-SDM-1-FP | | TGTGGTGGAATTCTGCAGATGAGCTCCAATGCGTCAACCTCACAAC | 2 kb |
| | | | SDM-3-FP | 3 nt insertion | TCTGCAGTCCTACCATGGATTTCAACCTACCAACGGA | |
| | | | SDM-4-RP | | GGTAGGTTGAAATCCATGGTAGGACTGCAGAGGGAAATAG | |
| | | | BEC-SDM-2-RP | | CGGCCGCCACTGTGCTGGATCTCGAGGTCTTCTAGGGGAGTTGGTCTG | |
| 4 | Single aa insertion | SARS-CoV-2 complete spike | BEC-SDM-1-FP | | TGTGGTGGAATTCTGCAGATGAGCTCCAATGCGTCAACCTCACAAC | 4 kb |
| | | | SDM-3-FP | 3 nt insertion | TCTGCAGTCCTACCATGGATTTCAACCTACCAACGGA | |
| | | | SDM-4-RP | | GGTAGGTTGAAATCCATGGTAGGACTGCAGAGGGAAATAG | |
| | | | BEC-SDM-4-FP | | CGGCCGCCACTGTGCTGGATCTCGAGTCAATTGAAGTTCACACACTTGT | |
| 5 | Long Deletion | SARS-CoV-2 complete spike | BEC-SDM-1-FP | 600 nt deletion | TGTGGTGGAATTCTGCAGATGAGCTCCAATGCGTCAACCTCACAAC | 1.5 kb |
| | | | SDM-5-FP | | GAGAGCATTGTGAGATTTCCCTTCAATGGACTGACCGGAA | |
| | | | SDM-6-FP | | TTCCGGTCAGTCCATTGAAGGGAAATCTCACAATGCTCTC | |
| | | | BEC-SDM-4-FP | | CGGCCGCCACTGTGCTGGATCTCGAGTCAATTGAAGTTCACACACTTGT | |
| 6 | Short deletion | SARS CoV-2 S1 | BEC-SDM-1-FP | | TGTGGTGGAATTCTGCAGATGAGCTCCAATGCGTCAACCTCACAAC | 2 kb |
|  |  |  | SDM-7-FP | 6 nt deletion | CCCTTTCTGGGAGTCTAC------AAGAACAATAAGAGCTGG |  |
|  |  |  | SDM-8-FP |  | CCAGCTCTTATTGTTCTT------GTAGACTCCCAGAAAGGG |  |
| | | | BEC-SDM-2-FP | | CGGCCGCCACTGTGCTGGATCTCGAGTCAATTGAAGTTCACACACTTGT | |
| 7 | Multiple mutations-1 | SARS CoV-2 S1 | BEC-SDM-1-FP | | TGTGGTGGAATTCTGCAGATGAGCTCCAATGCGTCAACCTCACAAC | 2 kb |
| | | | SDM-9-FP | | ATGCTACGGAGTGTCCCCTACCAAACTGAATGACCTCTGCTTCACAAACGTCTAT | |
| | | | SDM-10-RP | 8 nt substitution | GGACACTCCGTAGCATTTAAAGGTGAAGAAAGGGGCAAGATTGTAGAGGA | |
| | | | BEC-SDM-2-FP | | CGGCCGCCACTGTGCTGGATCTCGAGTCAATTGAAGTTCACACACTTGT | |

| | | | | | | |
|---|---|---|---|---|---|---|
| 8 | Multiple mutations-2 | SARS CoV-2 S1 | BEC-SDM-1-FP | | TGTGGTGGAATTCTGCAGATGAGCTCCAATGCGTCAACCTCACAAC | 2 kb |
| | | | SDM-11-FP | 9 nt substitution | TAAGGTTGATGGCCCATCCATAGGTAGGACGAAAAGAGTAGGATTTCAGAGGGA | |
| | | | SDM-12-RP | | GGGCCATCAACCTTATAGAGTGGTCGTGCTCAGCTTCGAGCTGCTGCATG | |
| | | | BEC-SDM-2-FP | | CGGCCGCCACTGTGCTGGATCTCGAGTCAATTGAAGTTCACACACTTGT | |

| | | | | | | |
|---|---|---|---|---|---|---|
| **9** | Multiple mutations-3 | SARS CoV-2 S1 | BEC-SDM-1-FP | | TGTGGTGGAATTCTGCAGATGAGCTCCAATGCGTCAACCTCACAAC | 2 kb |
| | | | SDM-13-FP | 8 nt substitution | CCATTACAAGGCTTGTTGCCGGCTTGGTAAATCTCGGTGCTAATGTCCCTCT | |
| | | | SDM-14-RP | | AACAAGCCTTGTAATGGCGTCGCTGGCTTCAACTGCTATTTCCCTCTGAAATCCT | |
| | | | BEC-SDM-2-FP | | CGGCCGCCACTGTGCTGGATCTCGAGGTCTTCTAGGGGAGTTGGTCTG | |
| **10** | Multiple mutations at multiple locations | SARS CoV-2 S1 | BEC-SDM-1-FP | | **TGTGGTGGAATTCTGCAGATGAGCTCCAATGCGTCAACCTCACAAC** | 2 kb |
| | | | SDM-9-FP | 17 nt substitution# | **ATGCTACGGAGTGTCCCCTACCAAACTGAATGACCTCTGCTTCACAAACGTCTAT** | |
| | | | SDM-10-RP | | **GGACACTCCGTAGCATTTAAAGGTGAAGAAAGGGGCAAGATTGTAGAGGA** | |
| | | | SDM-11-FP | | **TAAGGTTGATGGCCCACTCCATAGGTAGGACGAAAAGAGTAGGATTTCAGAGGGA** | |
| | | | SDM-12-RP | | **GGGCCATCAACCTTATAGAGTGGTCGTGCTCAGCTTCGAGCTGCTGCATG** | |
| | | | BEC-SDM-2-FP | | **CGGCCGCCACTGTGCTGGATCTCGAGGTCTTCTAGGGGAGTTGGTCTG** | |
| **11** | Multiple mutations at multiple locations | SARS CoV-2 S1 | BEC-SDM-1-FP | | **TGTGGTGGAATTCTGCAGATGAGCTCCAATGCGTCAACCTCACAAC** | 2 kb |
| | | | SDM-11-FP | 9 nt substitution* | **TAAGGTTGATGGCCCACTCCATAGGTAGGACGAAAAGAGTAGGATTTCAGAGGGA** | |
| | | | SDM-12-RP | | **GGGCCATCAACCTTATAGAGTGGTCGTGCTCAGCTTCGAGCTGCTGCATG** | |
|  |  |  | SDM-7-FP | 6 nt deletion** | **CCCTTTCTGGGAGTCTAC------AAGAACAATAAGAGCTGG** |  |
|  |  |  | SDM-8-FP |  | **CCAGCTCTTATTGTTCTT------GTAGACTCCCAGAAAGGG** |  |
| | | | BEC-SDM-2-FP | | **CGGCCGCCACTGTGCTGGATCTCGAGGTCTTCTAGGGGAGTTGGTCTG** | |

**#** S371L, S373P, S375F at one site; Q493K, G496S, Q498R, N501Y, Y505H at another site in the RBD
*Q493K, G496S, Q498R, N501Y, Y505H at RBD
** ΔY145 ΔH146 at NTD
**Red:** Mutation(s), **Blue:** Overlapping region, **Black:** Template-specific region, **Green:** Vector-specific region

**Table S5. List of overlapping primers used for synthesising short gene fragments using oligonucleotides**

| S. No | Oligo Code | Oligo Sequence (5' - 3') | GC % |
|---|---|---|---|
| 1 | OGS-BEC-1-FP | AGTGTGGTGGAATTCTGCAGATGAGCTCGCCACCATGAACATTACCAATCTGTGCCC | 50.9 |
| 2 | OGS-1-FP | AACATTACCAATCTGTGCCCCTTCGATGAGGTCTTTAACGCCACCAGATTCGC | 53.1 |
| 3 | OGS-2-RP | CGCAATTGGAGATCCTCTTTCTGTTCCAAGCATACACGGAAGCGAATCTGGTGGCG | 49.1 |
| 4 | OGS-3-FP | GGATCTCCAATTGCGTCGCCGACTACAGCGTCCTCTACAATCTTGCCCCT | 51.8 |
| 5 | OGS-4-RP | GGACACTCCGTAGCATTTAAAGGTGAAGAAAGGGGCAAGATTGTAGAGGA | 56 |
| 6 | OGS-5-FP | ATGCTACGGAGTGTCCCCTACCAAACTGAATGACCTCTGCTTCACAAACGTCTAT | 46 |
| 7 | OGS-6-RP | TGTCTCACTTCGTCGCCTCTGATGACAAAGCTATCGGCATAGACGTTTGTGAAGC | 47.3 |
| 8 | OGS-7-FP | CGACGAAGTGAGACAGATTGCCCCCGGACAGACCGGCAACATCGCCGATTACAAT | 49.1 |
| 9 | OGS-8-RP | TCACGCAGCCGGTAAAGTCGTCGGGCAGTTTGTAATTGTAATCGGCGATGT | 56.4 |
| 10 | OGS-9-FP | TTACCGGCTGCGTGATTGCTTGGAATAGCAACAAACTCGACAGCAAGGTGTC | 51 |
| 11 | OGS-10-RP | CTAAAGAGCCTATACAGATAGTTATAATTGCCAGACACCTTGCTGTCGAGT | 50 |
| 12 | OGS-11-FP | CTGTATAGGCTCTTTAGAAAGTCCAATCTGAAGCCCTTCGAGAGGGACATTAGCA | 41.2 |
| 13 | OGS-12-RP | CCATTACAAGGCTTGTTGCCGGCTTGGTAAATCTCGGTGCTAATGTCCCTCT | 45.5 |
| 14 | OGS-13-FP | AACAAGCCTTGTAATGGCGTCGCTGGCTTCAACTGCTATTTCCCTCTGAAATCCT | 50 |
| 15 | OGS-14-RP | TAAGGTTGATGGCCCACTCCATAGGTAGGACGAAAAGAGTAGGATTTCAGAGGGA | 47.3 |
| 16 | OGS-15-FP | GGGCCATCAACCTTATAGAGTGGTCGTGCTCAGCTTCGAGCTGCTGCATG | 47.3 |
| 17 | OGS-16RP | TGTGCTTTTCTTGGGTCCGCACACGGTGGCGGGAGCATGCAGCAGCTCGA | 56 |
| 18 | OGS-17-FP | CCCAAGAAAAGCACAAATCTGGTCAAGAACAAGTGTGTGAACTTCAAT | 62 |
| 19 | OSG-BEC-2-RP | CGGCCGCCACTGTGCTGGATCTCGAGTCAATTGAAGTTCACACACTTGT | 39.6 |

### 2. Supplementary figures

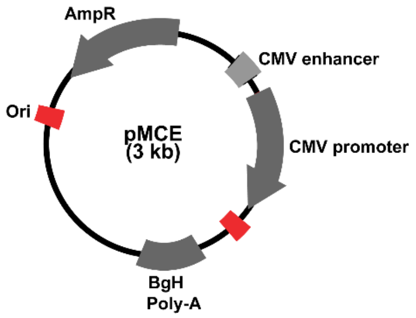


**Fig. S1. Plasmid map of pMCE eukaryotic expression vectors.** The plasmid pMCE derived from pcDNA 3.1 (+) consists of basic regulatory elements including origin of replication (ori), ampicillin resistance marker (Amp$^R$), CMV promoter, enhancer and BGH- polyadenylation signal.

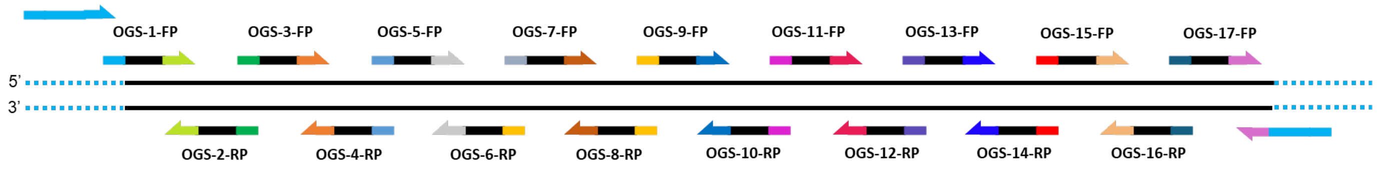


**Fig. S2. Schematic representation of the assembly of oligonucleotides used for constructing SARS-CoV-2 receptor binding domain (RBD).** OGS stands for oligonucleotide mediated gene synthesis.

### 3. Nucleotide sequence

#### pMCE vector

GCAAAAGGCCAGGAACCGTAAAAAGGCCGCGTTGCTGGCGTTTTTCCATAGGCTCCGCCCCCCTGACGAGCA
TCACAAAAATCGACGCTCAAGTCAGAGGTGGCGAAACCCGACAGGACTATAAAGATACCAGGCGTTTCCCCC
TGGAAGCTCCCTCGTGCGCTCTCCTGTTCCGACCCTGCCGCTTACCGGATACCTGTCCGCCTTTCTCCCTTC
GGGAAGCGTGGCGCTTTCTCATAGCTCACGCTGTAGGTATCTCAGTTCGGTGTAGGTCGTTCGCTCCAAGCT
GGGCTGTGTGCACGAACCCCCCGTTCAGCCCGACCGCTGCGCCTTATCCGGTAACTATCGTCTTGAGTCCAA
CCCGGTAAGACACGACTTATCGCCACTGGCAGCAGCCACTGGTAACAGGATTAGCAGAGCGAGGTATGTAGG
CGGTGCTACAGAGTTCTTGAAGTGGTGGCCTAACTACGGCTACACTAGAAGAACAGTATTTGGTATCTGCGC
TCTGCTGAAGCCAGTTACCTTCGGAAAAAGAGTTGGTAGCTCTTGATCCGGCAAACAAACCACCGCTGGTAG
CGGTTTTTTTGTTTGCAAGCAGCAGATTACGCGCAGAAAAAAAGGATCTCAAGAAGATCCTTTGATCTTTTC
TACGGGGTCTGACGCTCAGTGGAACGAAAACTCACGTTAAGGGATTTTGGTCATGAGATTATCAAAAAGGAT
CTTCACCTAGATCCTTTTAAATTAAAAATGAAGTTTTAAATCAATCTAAAGTATATATGAGTAAACTTGGTC
TGACAGTTACCAATGCTTAATCAGTGAGGCACCTATCTCAGCGATCTGTCTATTTCGTTCATCCATAGTTGC
CTGACTCCCCGTCGTGTAGATAACTACGATACGGGAGGGCTTACCATCTGGCCCCAGTGCTGCAATGATACC
GCGAGACCCACGCTCACCGGCTCCAGATTTATCAGCAATAAACCAGCCAGCCGGAAGGGCCGAGCGCAGAAG
TGGTCCTGCAACTTTATCCGCCTCCATCCAGTCTATTAATTGTTGCCGGGAAGCTAGAGTAAGTAGTTCGCC
AGTTAATAGTTTGCGCAACGTTGTTGCCATTGCTACAGGCATCGTGGTGTCACGCTCGTCGTTTGGTATGGC
TTCATTCAGCTCCGGTTCCCAACGATCAAGGCGAGTTACATGATCCCCCATGTTGTGCAAAAAAGCGGTTAG
CTCCTTCGGTCCTCCGATCGTTGTCAGAAGTAAGTTGGCCGCAGTGTTATCACTCATGGTTATGGCAGCACT
GCATAATTCTCTTACTGTCATGCCATCCGTAAGATGCTTTTCTGTGACTGGTGAGTACTCAACCAAGTCATT
CTGAGAATAGTGTATGCGGCGACCGAGTTGCTCTTGCCCGGCGTCAATACGGGATAATACCGCGCCACATAG
CAGAACTTTAAAAGTGCTCATCATTGGAAAACGTTCTTCGGGGCGAAAACTCTCAAGGATCTTACCGCTGTT
GAGATCCAGTTCGATGTAACCCACTCGTGCACCCAACTGATCTTCAGCATCTTTTACTTTCACCAGCGTTTC
TGGGTGAGCAAAAACAGGAAGGCAAAATGCCGCAAAAAAGGGAATAAGGGCGACACGGAAATGTTGAATACT
CATACTCTTCCTTTTTCAATATTATTGAAGCATTTATCAGGGTTATTGTCTCATGAGCGGATACATATTTGA
ATGTATTTAGAAAAATAAACAAATAGGGGTTCCGCGCACATTTCCCCGAAAAGTGCCACCTGACGTCGACGG
ATCGGGAGATCTCCCGATCCCCTATGGTGCACTCTCAGTACAATCTGCTCTGATGCCGCATAGTTAAGCCAG
TATCTGCTCCCTGCTTGTGTGTTGGAGGTCGCTGAGTAGTGCGCGAGCAAAATTTAAGCTACAACAAGGCAA
GGCTTGACCGACAATTGCATGAAGAATCTGCTTAGGGTTAGGCGTTTTGCGCTGCTTCGCGATGTACGGGCC
AGATATACGCGTTGACATTGATTATTGACTAGTTATTAATAGTAATCAATTACGGGGTCATTAGTTCATAGC
CCATATATGGAGTTCCGCGTTACATAACTTACGGTAAATGGCCCGCCTGGCTGACCGCCCAACGACCCCCGC
CCATTGACGTCAATAATGACGTATGTTCCCATAGTAACGCCAATAGGGACTTTCCATTGACGTCAATGGGTG
GAGTATTTACGGTAAACTGCCCACTTGGCAGTACATCAAGTGTATCATATGCCAAGTACGCCCCCTATTGAC
GTCAATGACGGTAAATGGCCCGCCTGGCATTATGCCCAGTACATGACCTTATGGGACTTTCCTACTTGGCAG
TACATCTACGTATTAGTCATCGCTATTACCATGGTGATGCGGTTTTGGCAGTACATCAATGGGCGTGGATAG
CGGTTTGACTCACGGGGATTTCCAAGTCTCCACCCCATTGACGTCAATGGGAGTTTGTTTTGGCACCAAAAT
CAACGGGACTTTCCAAAATGTCGTAACAACTCCGCCCCATTGACGCAAATGGGCGGTAGGCGTGTACGGTGG
GAGGTCTATATAAGCAGAGCTCTCTGGCTAACTAGAGAACCCACTGCTTACTGGCTTATCGAAATTAATACG
ACTCACTATAGGGAGACCCAAGCTGGCTAGCGTTTAAACTTAAGCTTGGTACCGAGCTCGGATCCACTAGTC
CAGTGTGGTGGAATTCTGCAGATATCCAGCACAGTGGCGGCCGCTCGAGTCTAGAGGGCCCGTTTAAACCCG
CTGATCAGCCTCGACTGTGCCTTCTAGTTGCCAGCCATCTGTTGTTTGCCCCTCCCCCGTGCCTTCCTTGAC
CCTGGAAGGTGCCACTCCCACTGTCCTTTCCTAATAAAATGAGGAAATTGCATCGCATTGTCTGAGTAGGTG
TCATTCTATTCTGGGGGGTGGGGTGGGGCAGGACAGCAAGGGGGAGGATTGGGAAGACAATAGCAGGCATGC
TGGGGATGCGGTGGGCTCTATGGCTTCTGAGGCGGAAAGAACCA
